# SLC12A9 is a lysosome-detoxifying ammonium – chloride co-transporter

**DOI:** 10.1101/2023.05.22.541801

**Authors:** Roni Levin-Konigsberg, Koushambi Mitra, AkshatKumar Nigam, Kaitlyn Spees, Pravin Hivare, Katherine Liu, Anshul Kundaje, Yamuna Krishnan, Michael C. Bassik

**Affiliations:** Department of Genetics, Stanford School of Medicine; Department of Chemistry, The University of Chicago; Neuroscience Institute, The University of Chicago; Institute of Biophysical Dynamics, The University of Chicago; Department of Computer Science, Stanford University; Department of Biology, Stanford University; Program in Chemistry, Engineering and Medicine for Human Health, Stanford University; Stanford Cancer Institute, Stanford School of Medicine

## Abstract

Ammonia is a ubiquitous, toxic by-product of cell metabolism. Its high membrane permeability and proton affinity causes ammonia to accumulate inside acidic lysosomes in its poorly membrane-permeant form: ammonium (NH_4_^+^). Ammonium buildup compromises lysosomal function, suggesting the existence of mechanisms that protect cells from ammonium toxicity. Here, we identified SLC12A9 as a lysosomal ammonium exporter that preserves lysosomal homeostasis. SLC12A9 knockout cells showed grossly enlarged lysosomes and elevated ammonium content. These phenotypes were reversed upon removal of the metabolic source of ammonium or dissipation of the lysosomal pH gradient. Lysosomal chloride increased in SLC12A9 knockout cells and chloride binding by SLC12A9 was required for ammonium transport. Our data indicate that SLC12A9 is a chloride-driven ammonium co-transporter that is central in an unappreciated, fundamental mechanism of lysosomal physiology that may have special relevance in tissues with elevated ammonia, such as tumors.

## Introduction

Ammonia is a ubiquitous metabolite that is produced by virtually all metazoan tissues and their associated microbiome. Maintaining low levels of systemic ammonia is critical because hyperammonemia can be toxic; high circulating ammonia causes neurotoxicity. Vertebrates prevent ammonia accumulation by converting it to urea ^1^ or recycling it for amino acid biosynthesis ^2^. However, these detoxification pathways are only active in a small subset of cell types and tissues ^1, 3^. Apart from the liver, which is the only organ that removes ammonia from circulation ^4^, little is known about how other tissues handle excess ammonia. Notably, abnormally high levels of ammonia have been reported in solid tumors ^5^, likely because of their accelerated metabolism and poor vascularization, suggesting that malignant cells have efficient mechanisms to counter this burden. Targeting such mechanisms may therefore be a useful strategy to curtail the growth of cancer cells in specific pathophysiological contexts.

Physiological ammonia (NH_3_) is in equilibrium with its protonated form, ammonium (NH_4_^+^), as its pKa is 9.01 ^6^. In serum (pH ≈ 7.4), free NH_3_ is estimated at 0.84 – 0.96 µM, nearly 40 times lower than NH_4_^+^ levels (**Fig. S1**). NH_3_ is neutral and hence, rapidly passes across most biological membranes. NH_3_ levels are therefore considered to be similar across all cellular compartments (**Fig. S1**). In contrast, NH_4_^+^ levels are dictated by the local luminal pH as it is charged and therefore cannot permeate membranes easily. Thus, NH_4_^+^ is predicted to substantially accumulate in highly acidic organelles such as lysosomes. Given a luminal pH of −4.5 and −0.84 –0.96 µM free NH_3_, the equilibrium levels of lysosomal NH_4_^+^ are estimated at 27 – 31 mM (**Fig S1**). This preferential and continual accumulation of NH_4_^+^ in lysosomes can conceivably affect their function, by altering their osmolarity, buffering power, and overall ionic balance. Indeed, that the extracellular addition of millimolar levels of NH_4_^+^ causes lysosomes to rapidly neutralize and stop functioning is well known ^7–11^. Thus, lysosomes must possess mechanisms to actively export NH_4_^+^ to the cytosol.

Here, we performed a genome-wide screen to identify regulators of endolysosomal traffic. In addition to known regulators of this pathway, we identified a critical new lysosomal ion transporter, SLC12A9. Using a combination of genetic, biochemical and microscopy approaches, as well as molecular dynamics simulations, we uncover SLC12A9 as the first putative lysosomal NH_4_^+^ – Cl^-^ cotransporter that has a critical role in the detoxification of ammonia generated metabolically. Additionally, we show that deletion of SLC12A9 causes a profound swelling of lysosomal compartments and sensitizes pancreatic ductal adenocarcinoma cells to NH_4_^+^ concentrations that have been reported to be found in solid tumors. Together, this work defines SLC12A9 as a critical regulator of lysosomal homeostasis.

## Results

### A CRISPR-Cas9 screen identifies SLC12A9 as a gene involved in endomembrane homeostasis

We began our study with the aim to uncover novel regulators of endocytic membrane traffic in macrophages. To this end, we installed a genome-wide sgRNA library in Cas9-expressing J774A.1 murine macrophages. Because we reasoned that abnormalities in endocytic traffic would result in changes in fluid phase uptake, we pulsed these cells with fluorescently-labeled dextran —a generic endocytic marker whose internalization is independent of cell surface receptors. These cells were then fixed and sorted based on the fluorescence intensity of the internalized dextran into two bins: top 10% (genes whose inactivation increased internalization) and bottom 12% (genes whose inactivation impaired uptake) (**Fig. 1A**). As expected, many of the hits that decreased uptake were genes related to actin cytoskeleton dynamics, which are known to be directly required for fluid phase uptake (macropinocytosis) ^12, 13^. We also found that numerous genes involved in endocytic trafficking such as RAB GTPases (RAB5C, RAB7, RAB1, RAB35), Rab-interacting proteins (ANKFY1, GAPVD1, WDR91, RIN2), phosphoinositide-related proteins (WDFY3, BECN1, UVRAG, PREX1, PIK3C3, INPP5F), membrane sorting and fusion machinery (SNX13, SNX24, VPS18, VPS11, VPS39), and microtubule transport motors (Dynein and Dynactin subunits) were hits in the screen (**Fig. 1B and C**), validating our experimental design.

**Figure 1.**
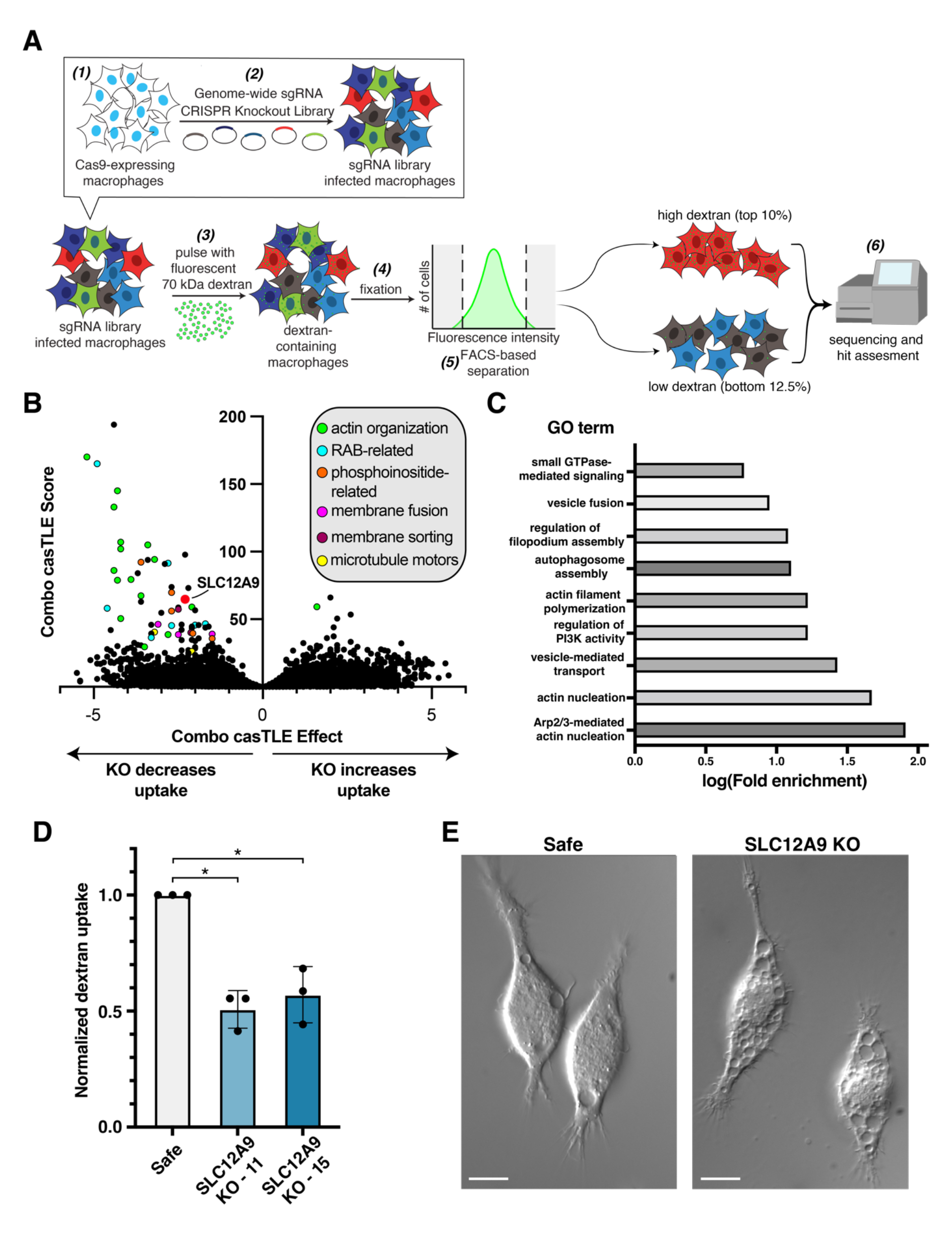
A genome-wide CRISPR/Cas9 screen to identify regulators of endocytic. trafficking uncovers SLC12A9 as a regulator of endomembrane volume regulation. **A.** Schematic showing the strategy for the genome-wide screen of regulators of macropinocytosis. **B.** Volcano plot of the screen depicted in **A. C.** Representative enriched cellular pathways (PantherDB, gene ontology term analysis) of the hits (141 genes within 10% FDR) from the screen. Redundant pathways (e.g., actin filament polymerization vs. actin polymerization; autophagosome assembly vs. macroautophagy) were consolidated into the annotated pathway with the higher enrichment score. **D.** Quantification of uptake of TMR-dextran by J774A.1 macrophages (Safe and SLC12A9 KO lines) using confocal microscopy. Shown are the means ± standard deviations of three independent experiments with at least 75 cells quantified per condition per replicate. Significance was determined using two-tailed unpaired t-tests **E.** Representative Nomarski micrographs of Cas9-J774A.1 cells transduced with a safe-targeting sgRNA (left) or with an SLC12A9-targeting sgRNA (right). Scale bars = 5 µm.

Among the hits with no previous association to endocytic trafficking we found SLC12A9 (**Fig. 1B**), an “orphan” member of the cation – chloride cotransporter (CCC) family ^14–16^ (**Fig. S2A**). To validate the involvement of SLC12A9 in endocytic traffic, we generated Cas9-expressing cell lines treated with individual sgRNAs targeting this gene. When we tested two SLC12A9 knockout (KO) J774A.1 lines (**Fig. S2B**), both showed reduced fluid phase uptake compared to control cells treated with sgRNA targeting safe regions within the genome (Safe) (**Fig. 1D**). Strikingly, however we also noticed that these KO cells accumulated abnormally large vacuoles throughout the cell (**Fig. 1E**). While WT J774A.1 macrophages commonly exhibit one or two transiently large, dynamic compartments, likely nascent macropinosomes, SLC12A9 KO cells exhibited multiple large, stationary structures that occupied a large fraction of the cell (**Fig. 1E**). Given this conspicuous observation, we sought to elucidate the molecular mechanisms behind this phenotype, which suggest a hitherto unappreciated function of SLC12A9.

### SLC12A9 localizes to lysosomes and is critical for their homeostasis

The SLC12A gene family encodes the well characterized cation – chloride co-transporters (CCC). Seven of the nine members of the family transport sodium and/or potassium along with chloride through biological membranes —mainly across the plasma membrane ^14, 16^ (**Fig. S2A**). In this family, SLC12A9, is an “orphan” transporter whose substrate specificity and physiological roles remain undefined ^14, 16^. Early studies did not detect cation transport via SLC12A9, leading to a suggested function as an ancillary CCC-interacting protein (CIP) ^17^.

SLC12A9 encodes a 12-transmembrane domain-containing integral membrane protein that was previously reported to localize to the plasma membrane ^17^. However, given the substantially enlarged vesicles associated with its inactivation, we analyzed the amino acid sequence of both human and mouse homologs. We identified two conserved lysosomal targeting motifs: a dileucine-based motif with a glutamic acid four residues upstream (EXXXLL), and a tyrosine-based motif with a hydrophobic residue (leucine) three amino acids downstream (YXXL) ^18^ (**Fig. 2A**). To test its subcellular localization, we expressed a fluorescently tagged version of SLC12A9 in HeLa and J774A.1 cells. Interestingly, SLC12A9-GFP was found as punctate structures both in the perinuclear region and dispersed throughout the cytosol, suggesting localization to vesicular endomembranes rather than to the plasma membrane (**Fig. 2B**). More specifically, SLC12A9-GFP showed high co-localization with the endo-lysosomal markers (LAMP1 and LAMP2) but not with markers of the trans-Golgi network (SYNT6) or early endosomes (RAB5) (**Fig. 2C and D**). To test whether the observed localization of SLC12A9-GFP was due to its native lysosomal localization sequences rather than an artifact of protein tagging and over-expression, we mutated the putative lysosomal-targeting motifs. Mutation of the dileucine motif (LL8/9AA) resulted in robust re-localization of SLC12A9-GFP to the plasma membrane, with a small fraction remaining in lysosomes (**Fig. 2E and F**). However, when we mutated both lysosomal-targeting motifs (LLY8/9/11AAA) we observed a complete mis-localization of SLC12A9 to the plasma membrane (**Fig 2E and F**). This indicates that SLC12A9 primarily localizes to lysosomal compartments.

**Figure 2.**
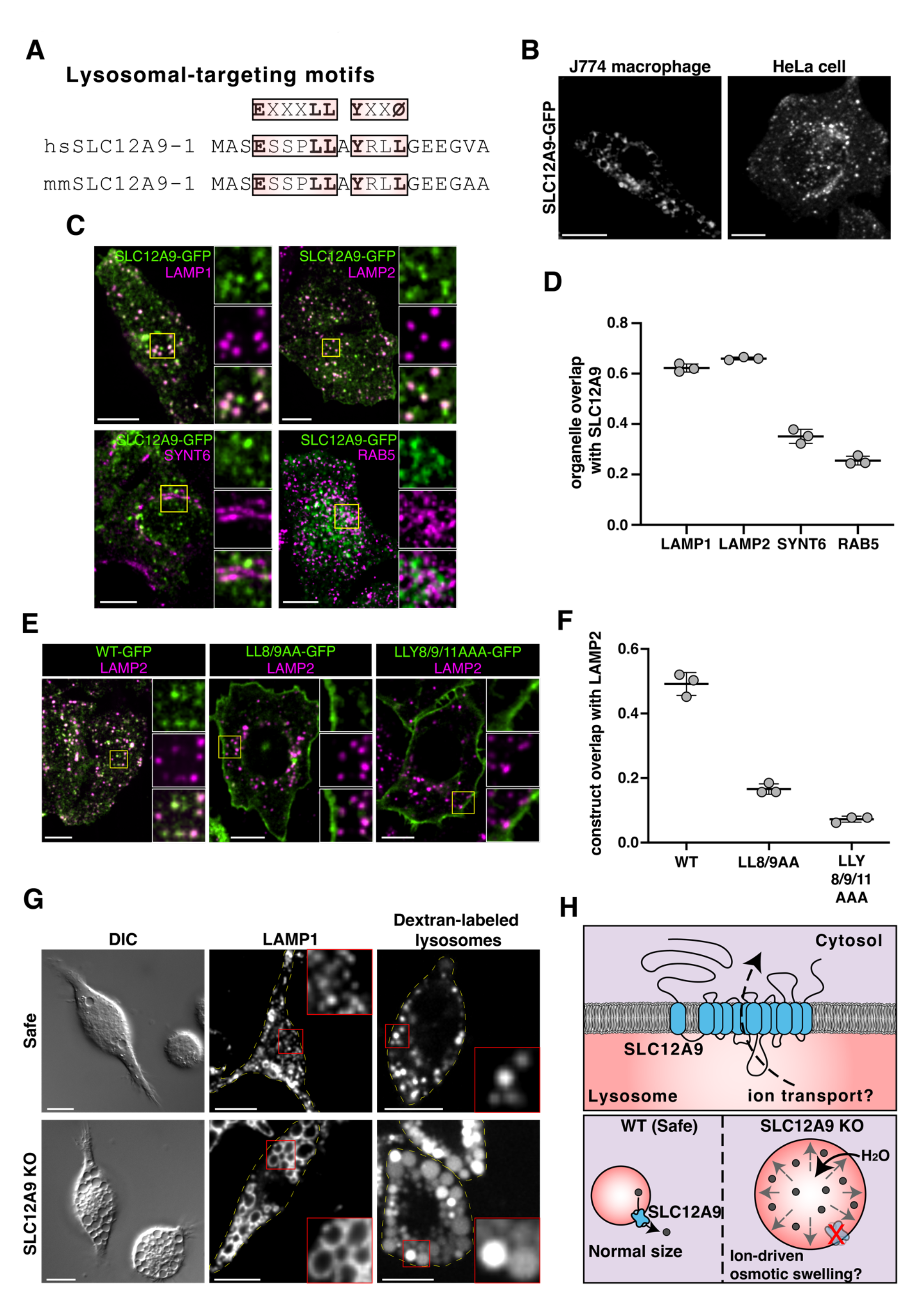
SLC12A9 localizes to and functions in lysosomes. **A.** Amino acid sequences of the N-terminal portion of SLC12A9 (human and mouse) highlighting lysosomal-targeting motifs are highlighted in boxes. **B.** Representative confocal slices of a J774A.1 macrophage (left) and a HeLa cell (right) expressing SLC12A9-GFP **C.** Representative confocal slices of HeLa cells expressing SLC12A9-GFP (green) and stained with antibodies (magenta) against: LAMP1 (endolysosomal marker), LAMP2 (lysosomal marker), SYNT6 (Golgi apparatus marker), and RAB5 (early endosomal marker). **D.** Co-localization analysis of indicated organelle markers with SLC12A9-GFP using Mander’s overlap coefficient (organelle overlap with SLC12A9-GFP to determine the proportion of each organelle that contains SLC12A9). Shown are the means and their standard deviations of three independent experiments with at least 50 cells quantified per condition per replicate. **E.** Representative confocal slices of HeLa cells stained with a LAMP2 antibody (magenta) and expressing GFP-tagged (green): SLC12A9-WT; SLC12A9-LL8/9AA; and SLC12A9-LLY8/9/11AAA. **F.** Co-localization analysis of SLC12A9-GFP variants with LAMP2 using Mander’s overlap coefficient (construct with LAMP2 to determine the proportion of the construct that localizes to LAMP2-positive compartments). Shown are the means ± standard deviations of three independent experiments with at least 40 cells quantified per condition per replicate. **G.** Representative micrographs of Cas9-J774A.1 murine macrophages expressing a safe-targeting sgRNA (top) or an SLC12A9-targeting sgRNA (bottom); Nomarski micrographs (left); confocal slices of cells stained with a LAMP1 antibody (center); confocal slices of cells pulsed with 10 KDa tetrametylrhodamine-dextran (2 h) and chased overnight (right). **H.** Schematic illustrating the hypothetical function of SLC12A9 as a lysosomal ion transporter (top) and the potential effects on lysosomal volume of its inactivation (bottom). Scale bars = 10 µm.

Given the lysosomal localization of SLC12A9-GFP, we tested whether the enlarged compartments observed in SLC12A9 KO cells (**Fig. 1E**) were lysosome-related. In this context, it is noteworthy that macrophages, which have a particularly rich and dynamic endolysosomal system, express high levels of SLC12A9 relative to most other cell types (**Fig S2C**). We found that these large compartments generated by SLC12A9 KO cells were at least partly of lysosomal origin, as the boundary of every swollen compartment had LAMP1 (**Fig 2G**). Further, lysosomal labeling with comparatively small fluorescent dextran (10 kDa) confirmed that the swollen compartments in SLC12A9 KO cells were indeed lysosomes (**Fig 2G**). Our data thus far suggested that SLC12A9 function is critical for lysosomal homeostasis.

### SLC12A9 transports NH_4_^+^ across biological membranes

Given that SLC12A9 is a member of the CCC (SLC12A) family of ion transporters, we reasoned that the lysosomal swelling caused by its deletion may be related to ion homeostasis (**Fig. 2H**). In addition, we noticed that the lysosomal phenotype was more pronounced in highly confluent cells that had not been passaged, suggesting a potential link with metabolism (**Fig S3A**). Interestingly when cells were washed and incubated in fresh medium or buffer (HBSS) the sizes of lysosomes were restored to normalcy over 1-2 hours (**Fig S3B and C**). This suggested that a product of cellular metabolism that had accumulated in the culture medium was inducing the lysosomal phenotype. Moreover, upon its removal from the medium, the metabolite was able to leave the lysosomes and restore their normal size. Interestingly, a recent screen found that cells lacking SLC12A9 are uniquely sensitive to the addition of ammonium chloride (NH_4_Cl). Notably, this effect could not be attributed to lysosomal alkalinization, as it was not phenocopied by V-ATPase inhibition ^19^. However, because the pair NH_3_/NH_4_^+^ is widely known as a product of cell metabolism, and because NH_3_ is membrane permeant, we reasoned that this couple fulfills all the criteria expected for the putative metabolite in our studies.

We therefore decided to test a model where NH_4_^+^ accumulates in lysosomes, due to their acidic luminal pH, when extracellular NH_3_ is elevated due to cell metabolism and triggers lysosome swelling in the absence of SLC12A9. When extracellular NH_3_ is removed, the high membrane permeability of NH_3_ coupled with the rapid conversion of lysosomal NH_4_^+^ to NH_3_ would rapidly reset the luminal levels of both species to normalcy (**Fig. S1**). Further, the ionic radius and charge of NH_4_^+^ are virtually identical to those of K^+ 20^, which is a substrate of most members of the SLC12A family of transporters ^14, 16^.

To test whether NH_4_^+^ accumulation was responsible for the phenotype of SLC12A9 KO cells, we assessed the effects of altering NH_3_/NH_4_^+^ on lysosome volume. In metazoans, glutamine metabolism is considered the main source of NH_3_^1^. We therefore compared lysosome size (determined by LAMP1 staining) as a function of cellular levels of NH_3_ + NH_4_^+^ (measured using a modified Berthelot reaction ^21^; **Fig. S3D**) with and without extracellular glutamine (**Fig. 3A**). Indeed, NH_3_ + NH_4_^+^ was present in control (safe) cells when cultured with glutamine but was barely detectable in its absence. However, the cellular content of NH_3_ + NH_4_^+^ increased massively (by 7 – 9-fold) in SLC12A9 KO cells incubated with glutamine (**Fig 3C**). Importantly, this accumulation was not observed in the absence of glutamine, implying that glutamine metabolism is the main source of cellular NH_3_ + NH_4_^+^ (**Fig 3C**).

**Figure 3.**
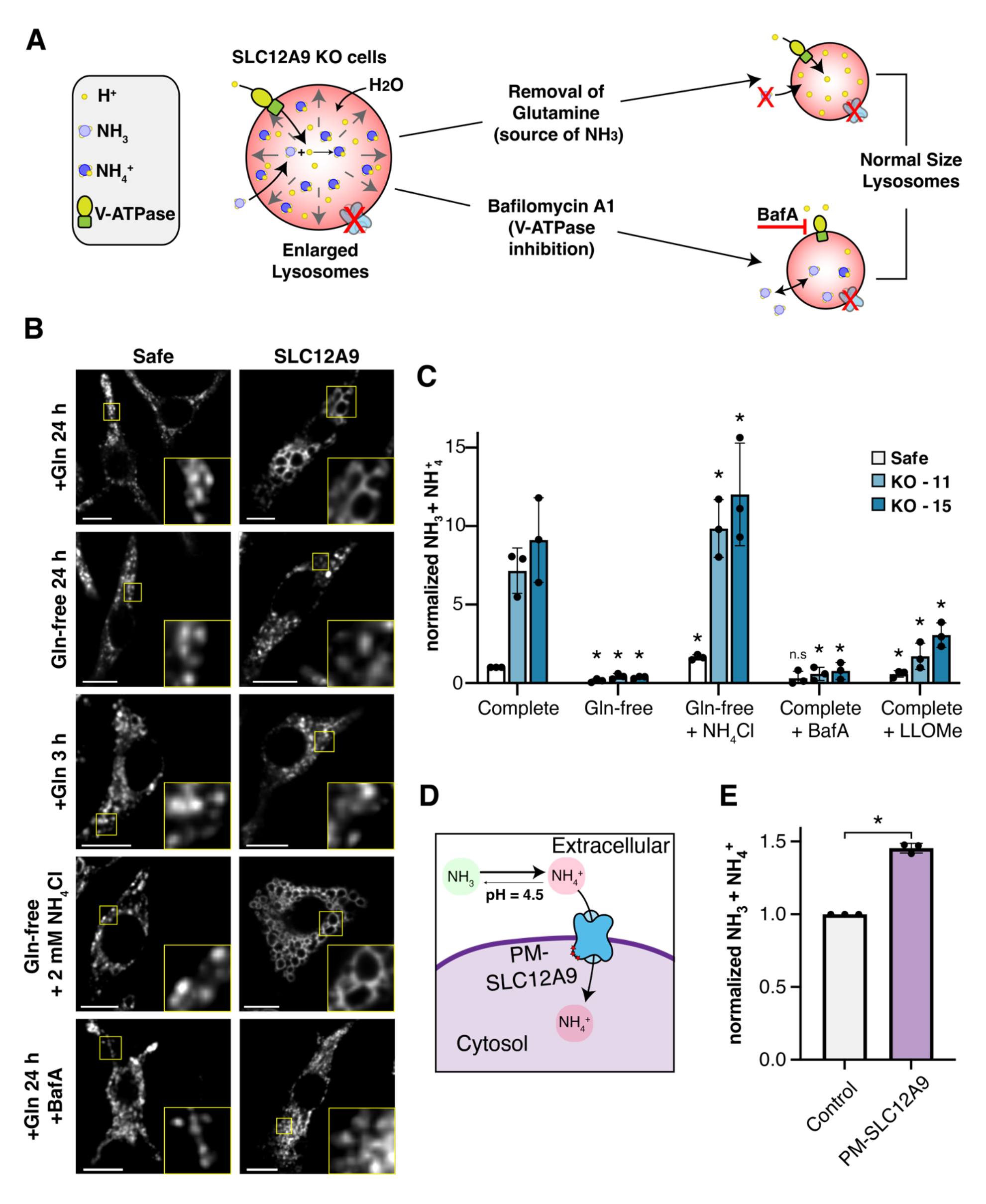
SLC12A9 transports ammonium across biological membranes. **A.** Schematic illustrating hypothetical conditions which would prevent ammonium-driven lysosomal swelling in SLC12A9 KO cells; top: removal of the metabolic source of ammonia: glutamine; right: dissipation of the lysosomal proton gradient: inhibition of the V-ATPase with Bafilomycin A. **B.** Representative confocal slices showing LAMP1 staining of control (top) and SLC12A9 KO (bottom) J774A.1 macrophages cultured under the described conditions. **C.** Quantification of total cellular ammonia + ammonium of control and SLC12A9 KO J774A.1 lines under the specified culture conditions using a modified version of Berthelot’s reaction. Shown are the means ± standard deviations of three independent experiments. Significance was calculated by two-tailed unpaired t-tests each cell line individually comparing culture in complete (control) medium and each of the alternative growth conditions; \**p* < .05 **D.** Schematic illustrating the ammonium transfer model at the plasma membrane using the plasma-membrane-localized SLC12A9-LLY8/9/11AAA mutant. **E.** Quantification of cellular ammonia + ammonium in control and PM-SLC12A9-transduced J774A.1 cells pulsed with acidic (pH = 4.5) ammonium-containing medium. Shown are means ± standard deviations of three independent experiments. Significance was calculated by a two-tailed unpaired t-test \**p* < .05. Scale bars = 10 µm.

Several lines of evidence suggest that NH_4_^+^ accumulation in lysosomes induces lysosomal swelling in SLC12A9 KO cells. First, omitting glutamine completely abolished swelling (**Fig 3B**). To test whether NH_4_^+^ was responsible for the swelling (rather than glutamine), we first cultured cells in glutamine-free medium, which prevented lysosome enlargement. Addition of glutamine for 3 h, a period deemed sufficient for glutamine uptake and equilibration in lysosomes, but insufficient for it to be metabolized to form significant amounts of NH_3_ / NH_4_^+^ to equilibrate to high levels in the extracellular medium, the cytosol and lysosomes, did not enlarge lysosomes in SLC12A9 KO cells (**Figs. 3B**). This suggests that a metabolic product of glutamine, and not glutamine itself, was responsible for the phenotype. In contrast, exogenous addition of NH_4_Cl to cells in glutamine-free medium induced a pronounced enlargement of lysosomes in SLC12A9 KO cells, while those of control cells were unaffected (**Fig. 3B**). Moreover, while cellular levels of NH_3_ + NH_4_^+^ levels increased markedly in KO cells treated with exogenous NH_4_^+^, those of control cells were several-fold lower (**Fig. 3C**).

Since NH_4_^+^ is predicted to accumulate in lysosomes because of their acidity, we tested whether dissipation of the lysosomal proton gradient would reduce NH_4_^+^ accumulation in SLC12A9 KO cells (**Fig. 3A**). When we inhibited the vacuolar ATPase (V-ATPase) with bafilomycin A1 (BafA) in SLC12A9 KO cells grown with glutamine, we found that NH_4_^+^ levels indeed diminished (**Fig. 3C**) and lysosomes did not swell (**Fig. 3B**), lending credence to the notion that swelling is a consequence of a pH-dependent NH_4_^+^ accumulation. Moreover, V-ATPase inhibition also prevented the lysosomal swelling in SLC12A9 KO cells subjected to exogenous addition of NH_4_Cl (**Fig. S3E**). Finally, disrupting the lysosomal membrane with L-leucyl-L-leucine methyl ester (LLOMe) reversed the accumulation of NH_3_ + NH_4_^+^ in SLC12A9 KO cells (**Fig. 3C**) indicating that the NH_4_^+^ build-up occurred in the lysosomal lumen. Together, our data suggest that SLC12A9 regulates luminal levels of NH_4_^+^ in lysosomes, potentially by acting as a transporter.

To directly test whether SLC12A9 can transport NH_4_^+^, we leveraged the triple mutant SLC12A9 (LLY8/9/11/AAA) which mislocalized to the plasma membrane (PM-SLC12A9; **Figs. 2E and F and S3F**). Given the topology of SLC12A9 at the plasma membrane, we manipulated the extracellular medium and tested whether PM-SLC12A9 directly imports NH_4_^+^ into the cell (**Fig. 3D**). Cells expressing the PM-SLC12A9 were briefly pulsed with 5 mM NH_4_^+^ and total cellular NH_3_ + NH_4_^+^ levels were then measured. The pH of the extracellular solution was lowered to 4.5 to resemble the lysosomal luminal pH and to minimize the availability of the more permeant NH_3_, which will inevitably enter the cells and cause SLC12A9-independent formation of intracellular NH_4_^+^. We observed that the NH_3_ + NH_4_^+^ content of cells expressing the PM-localized version of SLC12A9 significantly exceeded that of control cells (an - 40% increase) (**Fig. 3E**), indicating that SLC12A9 can transport NH_4_^+^ across biological membranes.

### SLC12A9 is an NH_4_^+^ – Cl^-^ co-transporter

Since the CCC (SLC12A) family of transporters use chloride (Cl^-^) to translocate cations^14, 16^, we tested whether SLC12A9 transported NH_4_^+^ in a chloride-dependent manner. Notably, the amino acid residues that are critical for Cl^-^ transport in the extensively characterized transporter NKCC1 (SLC12A2) ^22, 23^ are conserved among members of the SLC12A family (**Fig. 4A**). We assessed the importance of these residues, by transfecting SLC12A9 KO cells with either wildtype SLC12A9 or with mutants predicted not to bind Cl^-^ (Y313A and Y429A) and analyzed the ammonium-induced phenotypes. Notably, when KO cells expressing SLC12A9 mutants (Y313A and Y429A) were cultured in medium containing 2 mM NH_4_Cl, reduced growth and swollen lysosomes were still observed, sharply contrasting with cells expressing WT-SLC12A9, which rescued both phenotypes (**Fig. 4B and C**). These observations indicate that the putative chloride-binding sites in SLC12A9 are critical for NH_4_^+^ transport.

**Figure 4.**
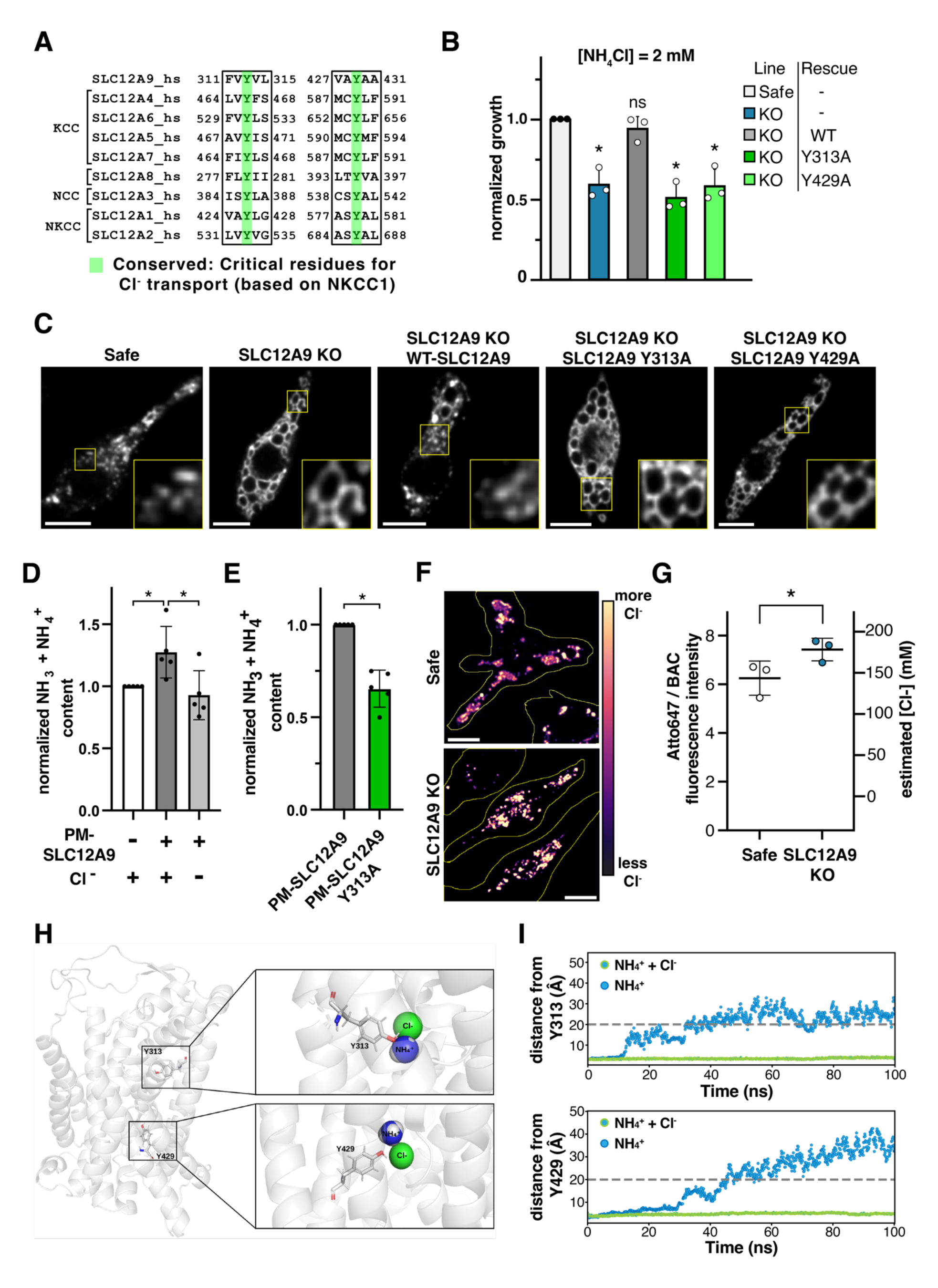
SLC12A9-mediated ammonium transport is chloride-dependent. **A.** Partial sequences of members of the SLC12A (CCC) family, highlighting conserved tyrosine residues that are critical for chloride transport by SLC12A2 (NKCC1). **B.** Quantification of the normalized growth during 72 h. in the presence of 2 mM NH4Cl of Safe or SLC12A9 KO J774A.1 cells expressing the listed rescue constructs. Shown are the means ± standard deviations of three independent experiments. Significance was calculated by two-tailed unpaired t-tests using the Safe line as a control. **C.** Representative confocal slices showing LAMP1 staining of the indicated J774A.1 cell lines, expressing the indicated rescue constructs. **D** and **E.** Quantification of total cellular ammonia + ammonium levels of: **D.** Control J774A.1 cells expressing a plasmalemmal-localized SLC12A9 mutant (LLY 8/9/11 AAA) in the presence or absence of chloride in the medium. Shown are the means ± standard deviations of five independent experiments. Significance was calculated by two-tailed unpaired t-tests. **E.** Control J774A.1 cells expressing the plasmalemmal-localized SLC12A9 mutant or its variant (Y313A) predicted to be incapable of binding chloride. Shown are the means ± standard deviations of five independent experiments. Significance was calculated by two-tailed unpaired t-tests. **F.** Representative pseudo-colored micrographs of Safe and SLC12A9 KO cells pulsed with the lysosomal chloride sensor Clensor - depicted is the ratio between Atto647 fluorescence and BAC. **G.** Quantification of lysosomal chloride levels shown as normalized ratiometric values, using Clensor in Safe and SLC12A9 KO cells. Shown are the means ± SD of three independent replicate experiments in which at least 90 lysosomes were quantified in at least 15 cells per experiment. Significance was calculated by two-tailed unpaired t-tests. **H.** AlphaFold predicted structure of the transmembrane core of SLC12A9; insets show still images of simulations of the predicted structure in the presence of ammonium and chloride, highlighting tyrosines (Y313 and Y429) predicted to be critical for chloride binding and ammonium transport. **I.** Quantification of the position of the ammonium ion relative to Y313 (top) and Y429 (bottom) during the above-mentioned simulations; dots with green outline: ammonium in the presence of chloride; dots with blue outline: ammonium in the absence of chloride. Scale bars = 10 µm.

We determined the need for Cl^-^ for SLC12A9 function by comparing the NH_4_^+^ transport capacity of PM-SLC12A9(**Figs. 2E and F and S3F**) in the presence or absence of extracellular Cl^-^. Interestingly, the cellular NH_3_ + NH_4_^+.^ levels of cells in chloride-containing buffers were significantly higher than those of cells incubated in chloride-free medium (−40% increase) (**Fig. 4D**). Accordingly, cells expressing the PM-localized transporter [SLC12A9 (LLY8/9/11/AAA)] with intact chloride-binding sites showed −30% higher NH_4_^+^ uptake than a PM-localized mutant in which the Cl^-^ binding residue Y313 is mutated (LLYY8/9/11/313AAAA) (**Fig. 4E, Fig. S4A and B**). This further implies that Cl^-^ binding by SLC12A9 is required for NH_4_^+^ transport.

Lysosomes are highly enriched in Cl^-^ largely due to the synergistic activity of V-ATPase and CLC-7, a chloride – proton antiporter ^24–26^. We therefore tested whether SLC12A9 is a chloride-activated NH_4_^+^ transporter or an NH_4_^+^ – Cl^-^ cotransporter. In the latter possibility, SLC12A9 inactivation would conceivably increase lysosomal Cl^-^. To assess this, we measured lysosomal Cl^-^ using Clensor ^25–27^, a ratiometric DNA-based Cl^-^ reporter which can be pulsed and chased into lysosomes of live cells (**Fig. S5 A – F**). We observed that lysosomes of SLC12A9 KO cells had a significantly higher concentration of Cl^-^ (an estimated excess of at least 20 mM) compared to those of control cells (**Fig. 4F – G and S5G – J**). These data suggest that SLC12A9 co-transports NH_4_^+^ and Cl^-^, conceivably using the lysosomal Cl^-^ gradient as a driving force for NH_4_^+^ translocation.

This model was further supported by molecular dynamics simulations in which we modeled the interactions between an AlphaFold generated structure of SLC12A9 (transmembrane core: residues 34 – 750) (**Fig. 4H**) ^28, 29^ with NH_4_^+^ and Cl^-^ ions placed proximally to Y313 and Y429. The simulations showed that the cation – anion pair was highly stable near these residues (**Fig. 4H - I, S4C** and **Supp. movie 1 - 2**). In contrast, in the absence of Cl^-^, the NH_4_^+^ ion was unstable and repelled from the transporter (**Fig. 4H - I,** and **Supp. movie 3 - 4**) consistent with our experimental data. Because it lacks a conserved tyrosine predicted to be critical for K^+^ transport in other CCC (SLC12A) family members (**Fig. S4E**), we hypothesized that SLC12A9 would not bind this cation, despite the fact that its ionic radius is very similar to that of NH_4_^+ 20^. We found that even in the presence of Cl^-^, K^+^ cannot be stabilized within the ion-binding pockets surrounding Y313 and Y429 of SLC12A9 (**Fig. S4F - G,** and **Supp. movie 5 - 6**) consistent with the idea that the transporter shows selectivity for NH_4_^+^ over K^+^. We cross-verified our simulations using AlphaFold predicted structures by comparing the results of similar modeling on NKCC1 (SLC12A2) with results obtained based on its CyroEM-determined structure ^22^. We separately modeled the pairs K^+^/Cl^-^ and Na^+^/Cl^-^ near Y533 and Y686, which are structurally equivalent to Y313 and Y429 in SLC12A9. Both cation – anion pairs were highly stable near Y533 and Y686, consistent with the original findings ^22^ and supporting our simulation results on SLC12A9 (**Figure S4H - I,** and **Supp. movies 7 - 10**).

### Physiological roles of SLC12A9

We next investigated the physiological role of SLC12A9 in lysosomal homeostasis. The excess NH_4_^+^ accumulation in lysosomes of SLC12A9 KO cells, suggested that their luminal pH could be consequently altered. Because deviations from the physiological pH are known to alter membrane traffic ^30–33^, an NH_4_^+^-driven alkalinization could therefore account for lysosomal swelling. However, lysosomal pH, measured using a ratiometric DNA-based pH reporter (I_m_^LY^) (**Fig. S6 A - G**) ^26^ showed no significant difference between control (safe) and SLC12A9 KO cells (**Fig. 5A - B**). Therefore, we speculated that lysosomal swelling was instead the direct result of the increased osmotic pressure associated with luminal NH_4_^+^ and Cl^-^ accumulation. Consistent with this, exposing SLC12A9 KO cells to a hyperosmotic solution caused swollen lysosomes to rapidly shrink and fragment to normal sizes, revealing that swelling was driven by osmotic pressure differences (**Fig. 5C**).

**Figure 5.**
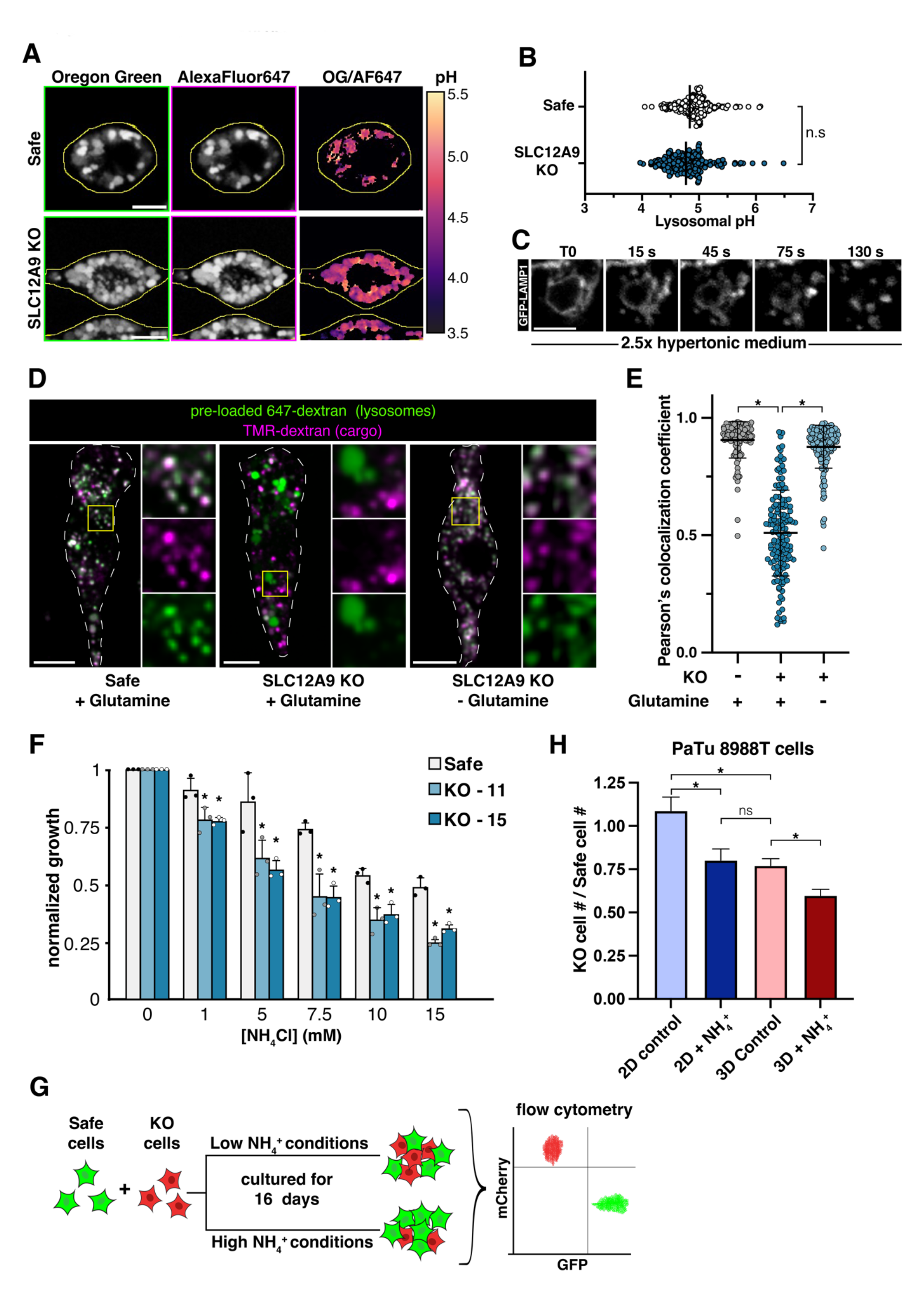
Physiological roles of SLC12A9. **A.** Representative confocal slices of safe and SLC12A9 KO J774A.1 cells pulsed with a DNA-based ratiometric (Oregon Green-488 - AlexaFluor-647) pH sensing probe. Rightmost panels show pseudo-colored micrographs depicting the ratio of fluorescence intensities of Oregon Green and AlexaFluor 647. **B.** Quantification of lysosomal pH of control and SLC12A9 KO J774A.1 macrophages based on the fluorescence intensity ratio illustrated in **A.** Shown are values of individual lysosomes and the means ± SD of the means of three independent replicate experiments in which at least 150 lysosomes were quantified in at least 10 cells per experiment. Significance was calculated by two-tailed unpaired t-tests. **C.** Representative time Lapse confocal slices showing the shrinkage of an enlarged lysosome of an SLC12A9 KO J774A.1 cell upon exposure to hyperosmotic (2.5x) medium. **D.** Representative confocal slices of the indicated J774A.1 lines with pre-labeled lysosomes (AlexaFluor 647 10 kDa dextran; green) subsequently pulsed with cargo (TMR 10 kDa dextran). **E.** Co-localization analysis of experiments shown in **D.** showing the Pearson’s colocalization coefficient between AlexaFluor 647 dextran and TMR dextran, each data point represents one cell. Shown are values of individual lysosomes and the means ± SD of the means of three independent replicate experiments. Significance was calculated by two-tailed unpaired t-tests. **F.** Normalized growth of Safe and SLC12A9 KO cell lines cultured in the presence of increasing concentrations of NH_4_Cl for 96 h. Each cell line’s growth was normalized to its confluency when cultured in the absence of ammonium. Shown are means + standard deviations of three independent experiments. Significance was calculated by two-tailed unpaired t-tests. **G.** Competitive growth assay for PaTu 8988T cells. Cells expressing negative control sgRNA (GFP) were mixed with an equal number of cells expressing SLC12A9-targeting sgRNA (mCherry) and grown either in 2D or 3D culture. Each culture type was grown in the presence or absence of ammonium chloride (2.5 mM). Shown are the means + standard deviations of the proportion of surviving SLC12A9 KO cells compared to the number of surviving Safe (control) cells under each culture condition from at least four independent experiments. Significance was calculated by two-tailed unpaired t-tests.

Deletion of SLC12A9 impaired fluid phase uptake (**Fig. 1D**), even though the co-transporter is largely restricted to lysosomes (**Fig. 2C - D**). This suggests that osmotic swelling of lysosomes could indirectly alter traffic along the entire endocytic pathway. Using tetramethylrhodamine (TAMRA)-dextran as an endocytic tracer, we tracked cargo delivery efficiency to terminal lysosomes pre-labeled with Alexa Fluor 647-dextran. SLC12A9 KO cells exhibited impaired delivery of the TAMRA-dextran to the lysosomes, in marked contrast to control cells (**Fig. 5D - E**). Importantly, this defect was rescued when SLC12A9 KO cells were incubated in glutamine-free medium (**Fig. 5D - E**). Thus NH_4_^+^ accumulation in lysosomes impaired endocytic trafficking.

We then tested whether altered lysosomal homeostasis due to NH_4_^+^ accumulation compromised cell viability. Addition NH_4_^+^ impaired the growth of SLC12A9 KO cells compared to control cells in a concentration-dependent manner (**Fig. 5F**), consistent with a recent report ^19^. In this regard, NH_4_^+^ has been reported to accumulate to remarkably high levels (1 – 5 mM) in solid tumors ^5^, which nevertheless grow robustly, indicating the significance of ammonia-detoxifying mechanisms in the tumor cells. Hence, we measured cell proliferation in culture environments with limited gas – liquid exchange that are predicted to promote NH_4_^+^ accumulation in the cell microenvironment. We knocked out SLC12A9 in PaTu 8988T cells —a pancreatic ductal adenocarcinoma cell line. Additionally, mCherry was expressed in these cells to aid in their identification. These KO cells were co-cultured with a safe (control) sgRNA-expressing PaTu 8988T line expressing GFP, and competitive growth assays in monolayer cultures and 3D spheroids ^34^ were performed. (**Fig. 5G**). While the SLC12A9 KO cells were not disadvantaged when grown in monolayers, they showed a significant proliferation impairment compared to safe cells when grown as 3D spheroids (**Fig. 5H**). As expected, addition of NH_4_Cl increased the sensitivity of SLC12A9 KO cells compared to safe cells in both 2D and 3D cultures (**Fig. 5H**). Notably, the growth disadvantage of SLC12A9 KO cells in spheroids cultured in regular medium was comparable to that of cells grown in 2D in the presence of NH_4_Cl (**Fig. 5H**). These data suggest that SLC12A9 may be critical for the survival of some cell types in solid tumors that are enriched in ammonia.

## Discussion

This work uncovers SLC12A9 as the first putative lysosomal NH_4_^+^ transporter, whose activity is critical for lysosomal homeostasis and function. Our data suggests that this protein operates as a critical NH_4_^+^ – Cl^-^ co-transporter that provides an efficient solution to the chemical problem arising due to the inherent tendency of physiological NH_3_ to enter the lysosomal lumen, get protonated and accumulate therein as NH_4_^+^, leading to lysosomal impairment.

SLC12A9, a hitherto “orphan” member of the CCC family, was previously thought to localize to the plasma membrane and to lack ion transport activity ^17^. Here we find instead that it is a lysosome-resident NH_4_^+^ exporter. SLC12A9 is critical for lysosomal homeostasis and function as its activity counters the continual permeation and protonation of NH_3_ in the acidic lysosome lumen. Given that the physiological extracellular levels of NH_3_/NH_4_^+^ are 35 – 40 µM, lysosomal NH_4_^+^ can theoretically approach −30 mM and impede lysosome function (**Fig. S1A**). Indeed, millimolar concentrations of NH_4_^+^ impair the activity of key lysosomal proteases such as cathepsin D ^35^. Moreover, NH_4_^+^ accumulation in lysosomes would elevate its osmolyte content and hydrostatic tension, both of which impair endomembrane traffic and lysosome reformation ^36–38^. Systemically, even comparatively modest increases in ammonia levels cause hyperammonemia. These observations highlight the significance of lysosomal NH_4_^+^ export (detoxification) mechanisms.

Overall, our data suggests that SLC12A9 functions as an NH_4_^+^ – Cl^-^ co-transporter. Osmolyte accumulation in lysosomes is promoted by the proton-driven import of Cl^-^ due to the activity of H^+^/Cl^-^ antiporter CLC-7, which has a 2Cl^-^/1H^+^ exchange stoichiometry ^24, 26, 39^. Cl^-^ acts as a neutralizing counterion that facilitates the continual electrogenic pumping of H^+^ into lysosomes. Additionally, Cl^-^ modulates cathepsin C activity in lysosomes ^40^. We show that Cl^-^ has an additional unappreciated role in lysosomal physiology by driving the export of NH_4_^+^ through SLC12A9, as the large lumen-to-cytosol Cl^-^ gradient likely provides the energy for NH_4_^+^ extrusion. By coupling the efflux of NH_4_^+^ to that of Cl^-^, lysosomes would efficiently counteract the osmotic burden imposed by the tendency of both ions to accumulate in their lumen. Remarkably, when correlating the expression of SLC12A9 to that of the entire proteome in a pan-cancer database ^41, 42^, CLC-7 expression has the sixth highest positive correlation. Interestingly, CLC-7 deletion in proximal tubular cells also leads to a robust lysosomal enlargement ^43^. While the mechanism behind this phenotype remains unknown, the predicted reduction in lysosomal Cl^-^ could impede NH_4_^+^ export resulting in an osmotically driven compartment enlargement.

Although SLC12A9 was previously thought to reside in the plasma membrane, we detected it primarily in lysosomes where its localization depended on two lysosomal-targeting motifs in its cytosolic portion. That SLC12A9 was previously thought to lack ion transport activity ^17^ may be attributable to the fact that NH_4_^+^ had not been tested as a possible substrate; optimal SLC12A9 function may also depend on the lysosomal environment. SLC12A9 has relatively low sequence similarity to other members of the CCC family ^14–16^ (**Fig. S2A**), supporting this unique substrate (NH_4_^+^) selectivity. Further, SLC12A9 lacks a critical tyrosine for K^+^ transport in NKCC1 ^22, 23^ (Y383 in the human homologue) that is conserved in K^+^ transporters of the CCC family. Accordingly, our molecular dynamics simulations predicted that K^+^ is unstable within the SLC12A9 ion-binding pockets surrounding Y313 and Y429, further supporting the selectivity of this transporter for NH_4_^+^, given similar charge and ionic radii of K^+^ and NH_4_^+ 20^.

SLC12A9 function is expected to be especially important in tissues enriched in NH_3_/NH_4_^+^. In this regard, SLC12A9 may be crucial in placental cells, where its expression is significantly higher than in other cell types (**Fig S2C**). Due to its high rates of metabolism, the placenta produces high levels of ammonia ^44–46^, suggesting that high SLC12A9 expression —and its NH_4_^+^ detoxifying function— are critical to protect placental cells from what otherwise would be a highly noxious environment during embryo development. In the context of pathology, the tumor microenvironment has been reported to be highly enriched in NH_3_/NH_4_^+^ (1 – 5 mM) ^5^ compared to normal circulating levels (≈40 μm). This is likely due to increased metabolism by cancer cells and limited liquid-gas exchange to and from tumors. Consistent with a greater role of SLC12A9 in such an environment, our data suggest that under the limited diffusion conditions inherent to 3D spheroids, SLC12A9 deletion results in a growth/survival disadvantage for cells. This raises the intriguing possibility of targeting SLC12A9 for therapeutic purposes, as it could potentially turn ammonia from a nutritional resource for cancer cells —which recycle ammonia for amino acid synthesis ^2^— into a toxic by-product.

## Methods

### Cell culture

All J774A.1 murine macrophage lines, HeLa cells and PaTu 8988t lines were cultured in DMEM (Gibco, 10313039) supplemented 10% fetal bovine serum, penicillin + streptomycin and L-glutamine at 37 °C with 5% CO_2_ unless specified otherwise for specific experiments.

### Endocytic traffic genome-wide CRISPR screen

Cas9-expressing J774A.1 murine macrophages were transduced sgRNA libraries (10 sgRNAs per gene + safe-targeting sgRNAs) and selected with puromycin (5 µg/ml). For the screen cells were used at 1500x coverage (1500 cells per guide within the library); cells were plated 24 h prior to exposure to 70-kDa FITC-dextran (Millipore Sigma, 46945). 3 h before the dextran treatment, cells were washed with D-PBS and incubated in serum-free medium. Then, cells were pulsed with 70 kDa FITC-dextran (25 µg/ml) in serum-free medium for 30 min. Next, cells were washed (2X) with D-PBS and (1X) with fresh serum-free medium. Media was then replaced with FBS-containing (5%) medium and cells were immediately spun-down. Cells were then resuspended in 4% paraformaldehyde (PFA) for fixation for 10 min, followed by a BSA-containing (5%) PBS wash. Fixed cells were then sorted based on FITC fluorescence intensity and binned into “top 10%” and “bottom 12.5%”. Genomic DNA from sorted cells and a control unsorted population was extracted and PCR-amplified in order to barcode the samples in preparation for NGS sequencing. Raw sequencing data was then demultiplexed and analyzed using CasTLE ^47^.

### Generation of cell lines

For generation of CRISPR-Cas9 KO lines, cells were transduced sgRNAs targeting SLC12A9 and incubated for 48 h for recovery. Then, cells were selected with puromycin (5 µg/ml for J774A.1 macrophages; 3 µg/ml for PaTu 8988t cells) for 24 h, followed by a 48 h incubation in regular medium for recovery and growth. For clonal lines, single cells were sorted into individual wells of 96-well plates and expanded.

### Dextran uptake assays

J774 murine macrophages were plated on glass coverslips 24 h prior to exposure to dextran. 3 h before dextran treatment, cells were washed with D-PBS and incubated in serum-free medium. Then, cells were pulsed with 70 kDa tetramethylrhodamine-labeled dextran (Thermo Fisher Scientific, D1818) (25 µg/ml) in serum-free medium for 30 min. Next, cells were washed (3X) with D-PBS and fixed with 4% PFA for 10 min.

### Lysosomal labeling in SLC12A9 KO cells

Cells were washed with PBS and incubated in glutamine-free medium for at least 2 h in order to restore lysosomal size and endocytic traffic. Cells were then pulsed with lysosomal markers or probes for varying times, washed and chased with a preconditioned medium (glutamine-containing medium from the original culture).

### Nomarski microscopy

Nomarski micrographs were obtained using an inverted Zeiss Axio Observer Z1 Epifluorescence / Widefield microscope equipped with an Orca Flash4.0 V3 sCMOS camara and an Axiocam 506 Color camera. Images were acquired using a Zen (Blue edition).

### Immunostaining

After each specific treatment, for all antibodies used except for anti-LAMP1 the cells were fixed with paraformaldehyde (4%) followed by a PBS wash. The cells were subsequently blocked with BSA (2%) for 40 min followed by a 1 h incubation with the primary antibody. The samples were then washed 3X with a BSA-containing (2%) solution and incubated with the fluorescent secondary antibody for 40 min in the dark. The samples were then washed with PBS (5X) and were subsequently imaged.

### Confocal Microscopy

Confocal micrographs were acquired using a Nikon Ti-E Eclipse base microscope equipped with a Yokogawa CSU-W1 scan unit and an ASI MS-2000 motorized X – Y stage. The system has a spectral unit with 405, 488, 561 and 640 nm laser lines. Images were acquired using an ANDOR iXon Ultra 987 EMCCD camera and a Nikon Plan Apo Lambda 60X oil objective, 1.4 NA, 0.15 mm WD. The system runs on NIS Elements v4 software.

● Nikon Ti-E Eclipse base
● Yokogawa CSU-W1 scan unit
● ASI MS-2000 motorized X-Y stage
● SPECTRAL laser unit (405, 488, 561, 640nm laser lines)
● ANDOR Zyla 4.2 scMOS camera
● ANDOR iXon Ultra 987 EMCCD camera
● Nikon Plan Apo Lambda objectives

10X air, 0.45NA, 4mm WD
20X air, 0.75NA, 1mm WD
40X air, 0.95NA, 0.21mm WD
60X oil, 1.4NA, 0.15mm WD
100X oil, 1.45NA, 0.13mm WD
● In Vivo Scientific incubation chamber

Images were analyzed using ImageJ (FIJI) version 2.1.0/1.53c

Co-localization and overlap were analyzed using the plug-in JACoP.

### Plasmids

The sgRNA library used in this study was cloned into pMCB320 (mCherry-tagged with Puromycin resistance cassette) and has been previously described ^48^. Individual sgRNAs for J774A.1 macrophages were cloned into pMCB307 (pMCB320 with BFP instead of mCherry). Individual sgRNAs for PaTu 8988t cells were cloned into pMCB320 (gene-targeting sgRNAs) or into pMCB406 (pMCB320 with GFP instead of mCherry; safe-targeting guides). sgRNA sequences used as follows: mmSLC12A9.1: GCAGGACTCTGGGGCCGG; mmSLC12A9.2: GGTTGCCTACATCATTC; mmSafe: GAAATGGATCTAACGCAGC; hsSLC12A9.1: GTCCTCTTTAACGGCTGTAC; hsSLC12A9.2: GCAGGACATAGACGAAGA; hsSafe1: GTTATAAAGTTAGATTTC; hsSafe2: GCCTACAGATGGTTACA.

SLC12A9-GFP was cloned by amplifying the ORF from pDONR221_SLC12A9 (Addgene plasmid #131897) and inserting it into a backbone containing GFP and a Blasticidin resistance cassette. Multicistronic constructs were designed using a mmSLC12A9 gene block from Twist Biosciences and inserting it into a backbone in which the gene was preceded by GFP-T2A and followed by P2A-Blasticidin resistance cassette. All site-directed mutant versions of SLC12A9 (i.e. LL8/9AA, LLY8/9/11AA, Y313A, Y429A and LLYY8/9/11/313AAAA) were generated through Gibson assembly.

### Antibodies

Antibodies used for this study:

Anti-LAMP1 (Developmental Studies Hybridoma Bank, ID4B); anti-LAMP1 (Developmental Studies Hybridoma Bank, H4A3); anti-LAMP2 (Developmental Studies Hybridoma Bank, H4A4); anti-Rab5 (BD Biosciences, 610724); anti-Synt6 (Cell Signaling, C34B2).

### Total cellular ammonia + ammonium measurement

Total cellular ammonia + ammonium was determined using a modified Berthelot reaction. Two solutions were prepared: Solution A contained 100 mM phenol (Millipore Sigma, P1037) and 50 mg/L sodium nitroprusside (Millipore Sigma, 71778); Solution B contained 0.38 dibasic sodium phosphate, 125 mM NaOH and 1% sodium hypochlorite (available chlorine 10 – 15%) (Millipore. Sigma, 425044).

On the day before the determination, 1.2 x 10^6^ cells were plated in 10-cm dishes. On the next day, cells were scrapped, centrifuged, and resuspended in preconditioned medium (medium from the original culture). Cells were then counted, and 1 x 10^6^ cells were centrifuged and resuspended in ice-cold HBSS. Cells were then briefly centrifuged and the supernatant was removed. The cell pellet was then lysed with 500 µl of ice-cold methanol:water (80:20) followed by a 10 minute incubation at −20 °C. Next, the lysates were spun-down at 10,000 g for 10 minutes at 4 °C. During the centrifugation 100 µl of Solution A were added to wells of 96-well plates. Lysates were then kept on ice and 20 µl of samples were added by triplicate into wells containing Solution A. The modified Bertheliot reaction was initiated by adding 100 µl of. Solution B into the wells. The plates were then incubated at 37 °C in the dark for 40 minutes. After this incubation the absorbance of each well was read at a wavelength of 635 nm using a plate reader.

For ammonium transport across the plasma membrane cells were scraped, washed and counted on the day of the experiment. 1.5 x 10^6^ cells were plated and incubated in FBS-containing medium. After 4 h., these cells were washed with D-PBS and incubated in glutamine-free medium for 4 h. Cells were then pulsed with a saline buffer (pH = 4.5) that contained 5 mM NH_4_Cl for 2 min at 37 °C. Next, cells were immediately washed 2X by dunking into sequential beakers containing ice-cold saline buffer (pH = 4.5) without ammonium. The cells were immediately lysed and total ammonium + ammonia was determined as described above. For chloride-free systems, the chloride containing salts in the saline buffer were replaced by nitrate-based salts (i.e., sodium nitrate, potassium nitrate, magnesium nitrate and calcium nitrate) and the ammonium-pulsing buffer contained ammonium gluconate.

### Multiple sequence alignment

The sequence analysis of the members of the SLC12A family (*Homo sapiens*) was performed using COBALT from the NCBI. Accession numbers used: SLC12A1: Q13621; SLC12A2: P55011; SLC12A3: P55017; SLC12A4: Q9UP95; SLC12A5: Q9H2X9; SLC12A6: Q9UHW9; SLC12A7: Q9Y666; SLC12A8: A0AV02; SLC12A9: Q9BXP2

### Phylogenetic homology analysis

The homology analysis for members of the SLC12A family was performed using the multiple sequence analysis described above, in the Phylogeny Analysis tool of *Methodes et Algorithmes pour la Bio-informatique*; LIRRM.

### DNA synthesis and characterization

The DNA sensors were synthesized and characterized using published protocols (Chakraborty *et al.*, eLife, 2017, 6:e28862). Briefly, equimolar amounts of desired ssDNA were dissolved in 10 mM sodium phosphate buffer (pH 7.4) and annealed by initial heating at 90°C for 5 min, followed by cooling at the rate of 5°C / 15 min. For preparation of 10 μM stocks of the pH sensor (*ImLy*) or Alexa647-labeled DNA duplex (*ImLy^mono^*), 10 μM of each *D2_ImLy_^Alexa647^* and *D1_ImLy_^OG488^* or *D1_ImLy_^unlabeled^* were annealed. Stock solutions of Clensor were similarly prepared at a final concentration of 10 μM by annealing equimolar ratio of *P1_Clensor_^Bac^*, *D1_Clensor_^Atto647^*, *D2* and *P* in 10 mM sodium phosphate buffer (pH 7.4).

### Reagents for pH and chloride measurements

All fluorescently labeled oligonucleotides (Table S1) were HPLC purified and purchased from Integrated DNA Technologies (IDT, Coralville, IA, USA). TMR conjugated 10kDa Dextran was obtained from Thermo Fisher Scientific. Nigericin, valinomycin, monensin, and tributyltin chloride were obtained from Sigma. The peptide nucleic acid (PNA) oligomer, P (Table S1) was synthesized using standard solid phase Fmoc chemistry on Nova Syn® TGA resin (Novabiochem, Germany) using analytical grade reagents (Applied Biosystems®, USA), purified by reverse phase HPLC (Shimadzu, Japan) and stored at −20 °C until further use. PNA-BAC was synthesized using protocols as described earlier (Prakash, Saha *et al.*, *Chem Sci.*, 2016, 7, 1946). All reagents for cell culture were purchased from Invitrogen Corporation, USA.

### *In vitro* bead calibration of *ImLy*

Briefly, bead calibration was performed using *ImLy* DNA labeled 1-μm streptavidin coated microspheres in 50 mM potassium phosphate buffer, pH 7.4, 150 mM NaCl with 0.05% Tween 20 for 2 h at room temperature. The beads were washed thrice by spinning at 12000 rpm for 10 mins each at room temperature. The beads adsorbed with *ImLy* were incubated with clamping buffers containing (HEPES (10 mM), MES (10 mM), sodium acetate (10 mM), KCl (140 mM), NaCl (5 mM), CaCl_2_ (1 mM) and MgCl_2_ (1 mM)) of desired pH (3.0, 3.5, 4.0, 4.5, 5.5, 5.5, 6.0, 6.5, and 7.0) for 30 mins at room temperature. After incubation, the bead containing solutions (2 μL) were imaged on a glass slide using IX83 inverted wide field microscope. Oregon green 488 (G), and Alexa 647 (R) was excited at 488 nm, and 647 nm respectively. G/R (pH) values, obtained from individual images from three independent experiments, were plotted for different points as function of pH.

### *In cellulo* pH clamping of *ImLy*

J774A.1 WT cells were labeled with 500 nM *ImLy* in serum-free DMEM media (containing 4.5 g/L glucose, sodium pyruvate and L-glutamine) for 1 h followed by a chase of 2 h in preconditioned imaging media. Cells were washed twice, fixed in 4% PFA in 1X PBS for 10 mins. After thorough washing, cells were clamped in clamping buffers (HEPES (10 mM), MES (10 mM), sodium acetate (10 mM), KCl (140 mM), NaCl (5 mM), CaCl_2_ (1 mM), MgCl_2_ (1 mM), nigericin (50 µM), and monensin (50 µM) for 1 h at room temperature. Cells were then imaged using IX83 inverted wide field microscope and ZEISS LSM 980 with Airyscan 2 confocal microscope.

### *In cellulo* pH measurements by *ImLy*

J774A.1 WT and SLC12A9 KO cells were pulsed with 500 nM *ImLy* in serum-free DMEM media (containing 4.5 g/L glucose, sodium pyruvate and L-glutamine) for 1 h followed by a chase of 2 h or in preconditioned imaging media. Cells were then imaged live by ZEISS LSM 980 with Airyscan 2 confocal microscope.

### *In cellulo* chloride clamping of *Clensor*

Chloride clamping and measurements in J774A.1 WT and SLC12A9 KO cells were carried out using *Clensor* following a previously published protocol (Saha *et al.*, Nature Nanotechnol., 2015, 10, 645–651, Chakraborty *et al.*, eLife, 2017, 6:e28862). J774A.1 WT cells were pulsed and chased with 2 µM of *Clensor*. Cells were then fixed with 100 µL of 2.5% PFA for 5 min at room temperature and washed three times with 1X PBS. After thorough washing, cells were then clamped for 1 h at room temperature in chloride clamping buffers containing a specific concentration of chloride, nigericin (50 µM), valinomycin (50 µM), and tributyltin chloride (50 µM). To prepare chloride clamping buffers of desired chloride concentrations, 1X chloride positive buffer (KCl (120 mM), NaCl (20 mM), CaCl_2_ (1 mM), MgCl_2_ (1 mM), HEPES (20 mM), pH, 7.2) and 1X chloride negative buffer (KNO_3_ (120 mM), NaNO_3_ (20 mM), Ca(NO_3_)_2_ (1 mM), Mg(NO_3_)_2_ (1 mM), HEPES (20 mM), pH 7.2) were mixed in the different ratios. Cells were then washed with 1X PBS and imaged using ZEISS LSM 980 with Airyscan 2 confocal microscope.

### *In cellulo* chloride measurement by *Clensor*

For real-time chloride measurements, J774A.1 WT and SLC12A9 KO cells were pulsed with 2 µM of *Clensor* for 1h followed by a 2 h or chase. Cells are then washed with 1X PBS and imaged live in preconditioned imaging media using ZEISS LSM 980 with Airyscan 2 confocal microscope.

### Fluorescence microscopy imaging for pH and chloride measurements

Bead calibration was done by using IX83 inverted wide field microscope (Olympus Corporation of the Americas) with 60x, 1.42 NA or 100X, 1.42 NA, differential interference contrast (DIC) objective (PLAPON, Olympus Corporation of the Americas) and Evolve Delta 512 EMCCD camera (Photometrics). For different fluorophores excitation and emission MetaMorph Premier Ver 7.8.12.0 software (Molecular Devices LLC, USA) was used for controlling filter wheel, shutter, and charge-coupled device camera. Oregon green 488 was imaged with 500/20 band-pass excitation filter, 520/40 band-pass emission filter and 89016 dichroic mirror. Alexa 647 was imaged with 640/30 band-pass excitation filter and 705/72 band-pass emission filter with 89016 dichroic. Images in Oregon green 488 channel were acquired with 1000 ms exposure time and 2000 EM Gain. Images in Alexa 647 channel were acquired with 400 ms exposure time and 1500 EM Gain. The confocal microscope used for these determinations is ZEISS LSM 980 with Airyscan 2 (Carl Zeiss) with Argon laser for 445 nm excitation and 488 nm excitation, Diode pumped solid state laser (DPSS) for 561 nm excitation, and HeNe laser for 639 nm laser excitation using Plan-Apochromat 63 X/1.4 oil DIC M27, 0.14 mm objective. BAC was excited with Argon laser at 445 nm and emission was collected from 452-540 nm; Oregon Green 488 was excited with Argon laser at 488 nm and emission was collected from 500-580 nm; TMR dextran was excited with DPSS laser at 561 nm and emission was collected from 570-640 nm; A647 was excited with HeNe laser at 639 nm and emission was collected from 650-760 nm. Images were acquired using GaAsP-PMT detectors. All images were acquired in sequential mode from both widefield and confocal microscopes.

### Image analysis

Images acquired using IX83 inverted wide field microscope and ZEISS LSM 980 with Airyscan 2 confocal microscope were analyzed with Fiji ImageJ ver 1.53q (NIH, USA).

### Colocalization

The colocalization of TMR dextran and Alexa 647 was determined by using Pearson’s correlation coefficient (PCC) using ImageJ. The PCC and pixel shift values for each time point were calculated using Coloc 2 test.

### pH measurements

For pH measurements Oregon Green 488 (G) and Alexa 647 (R) images were background subtracted, overlapped using ImageJ and lysosomes showing colocalization were selected for further analysis. The ROIs were applied to the images of G and R and the intensity was measured for each ROI. The ratio of G to R intensity (G/R) was then plotted for each pH point and the calibration curve was generated using Boltzmann function. The G/R values from *in cellulo* clamping and *in cellulo* pH measurements were converted to pH according to the *in vitro* bead calibration curve. Data was represented as the mean pH value ± standard error of the mean.

### Chloride measurements

For chloride measurements in J774A.1 WT and SLC12A9 KO cells, the lysosomes were identified in each Alexa 647 (R) image and the ROIs were marked using ImageJ for both Alexa 647 (R) and BAC (G) channels. The mean intensity was calculated for R and G after background subtraction and a ratio of R to G intensity (R/G) for each endosome was measured. The Cl values were determined from mean R/G values by using intracellular calibration profile. Data was presented as the mean Cl value ± standard error of the mean. Data for *in cellulo* chloride clamping experiments were analyzed similarly.

### Osmotic resolution of enlarged lysosomes

SLC12A9 KO J774A.1 cells were cultured on glass coverslips overnight in complete medium. The next day, the cells were washed with 1X short term imaging buffer (D-PBS + HEPES) + 50 µM NH_4_Cl and were placed in an imaging chamber with 500 µl of 1X buffer. After acquisition of the first micrograph at T0, 500 µl of 2X imaging buffer was added into the chamber and imaging was continuously acquired approximately every 15 s. depending on the stability of the focus of the previous micrograph.

### Molecular Dynamics Simulations

The predicted structure of SLC12A9 was obtained from https://alphafold.ebi.ac.uk/entry/Q9BXP2 and prepared using Schrödinger’s protein preparation wizard. For simulations, the protein was oriented with the Orientations of Proteins in Membranes (OPM) webserver v.2.0, solvated in a box of lipid bilayer, TIP3P water and 150mM KCl with the CHARMM-GUI. The Gromacs (v2021.3) software package was used to execute simulations using a Langevin thermostat and a Nosé– Hoover Langevin piston barostat at 1atm with a period of 50fs and decay of 25fs. The system was minimized for 5,000 steps, heated to 303.15K. This was followed by an equilibration, consisting of 125000 steps at a 1fs time step. Production simulations were performed for over 100ns, using the CHARMM36 force-field. All simulations were run in replicates of 5. The distances between ions and residues of all plots were calculated using the MDTraj python package. Visualization of the generated trajectories was done using PyMol.

### Competitive growth assays

Safe (GFP-expressing) and SLC12A9 KO (mCherry-expressing) PaTu 8988t cells were treated with accutase and counted separately using an Attune NxT cytometer. Next, cells were mixed in equal proportions and counted again, the resulting proportion between the cell lines was determined as T0. For 2D culture conditions, 100,000 cells per well were plated into 6-well plates; for 3D conditions, 100,000 cells per/well were plated into ultra-low attachment 6-well plates with methylcellulose-containing media ^34^. For each culture type (2D and 3D) a control medium and an NH_4_Cl-containing (3 mM) medium conditions were tested. Medium in the wells was replaced with corresponding fresh medium on days 2, 6, 10 and 14. Cells were split on days 4, 8, 12 and 16. Cell splitting for 3D cultures was performed as in ^34^. Cell counting was performed using an Attune NxT cytometer.

## Contributions

R.L.K and M.C.B conceived and designed the study. R.L.K performed and analyzed the macropinocytosis genome-wide screen. K.S cloned the sgRNA libraries. R.L.K and K.S cloned and validated the efficiency of the sgRNAs used in the study. R.L.K cloned the additional constructs used in this study. R.L.K performed the Nomarski, brightfield and confocal microscopy and their associated analyses with the exception of the lysosomal pH and chloride determination experiments, the macropinocytosis assays, the growth/kill curves upon ammonia exposure. R.L.K and K.S. performed the cellular ammonia + ammonium determinations. R.L.K performed the sequence conservation analysis. K.M synthesized Clensor and performed the analysis for the lysosomal pH and chloride determinations. P.H synthesized the pH probe (I_m_^Ly^), cultured cells, standardized the labeling conditions for ImLy and Clensor and acquired the images for these determinations A.N analyzed the AlohaFold predicted structures used in the study. A.N. performed and analyzed the molecular dynamics simulations with the advice of A.K. R.L.K performed and analyzed the lysosomal delivery assays. R.L.K and K.L. performed and analyzed the competitive growth assays with the help of K.S. K.L. optimized the 3D growth conditions for PaTu 8988T cells. R.L.K. analyzed the expression profiles of SLC12A9. R.L.K and M.C.B wrote the manuscript. All authors reviewed and edited the manuscript.

## Acknowledgements

We thank the Stanford Shared FACS Facility for the instruments and support for flow cytometry. We thank the Cell Sciences Imaging Facility at Stanford for instrumentation and support for Nomarski microscopy. We thank Dr. Monther Abu-Remaileh for the PaTu 8988t cells and for valuable discussions. We thank members of the Bassik lab for discussions, feedback and support. This work was supported by NCI U54CA261719 and NIH UM1HG011972 awarded to M.C.B; by R01 NS112139-01A1 and 1DP1OD033642-0 awarded to YK.

---

**Figure S1.**
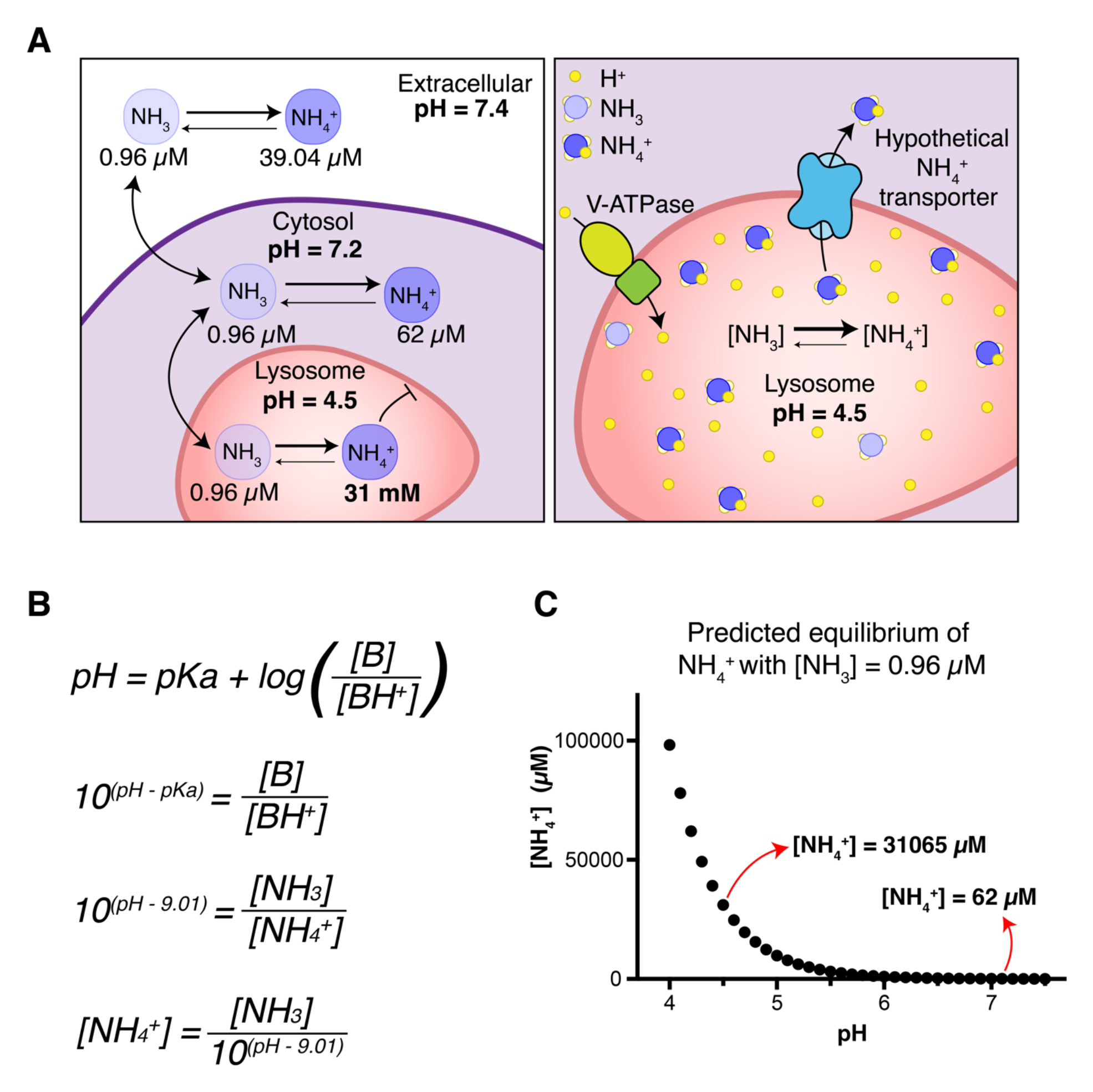
Ammonia – Ammonium equilibrium in cell physiology. **A.** Schematic showing: (left) the concentration of ammonia (NH_3_) —a highly membrane-permeant molecule— is assumed to be equal across cellular membranes; considering physiological levels of NH_3_ + NH_4_^+^ in circulation (40 µM) using the Henderson - Hasselbalch equation (**B.**) at extracellular pH (7.4), the concentration of ammonia is considered 0.96 µM across all cellular membranes. **C.** At extracellular and cytosolic pH (7.4 and 7.2 respectively) 0.96 µM NH_3_ is in equilibrium with 39.04 and 62 µM NH_4_^+^ respectively. At lysosomal pH (4.5) 0.96 µM NH_3_ is predicted to equilibrate with 31 mM NH_4_^+^. This tendency of NH_4_^+^ to accumulate in acidic compartments would conceivably have deleterious effects in lysosomal homeostasis and function, because of this we hypothesize that lysosomes have active mechanisms to prevent such a massive accumulation by exporting NH_4_^+^ to the cytosol (**A**).

**Figure S2.**
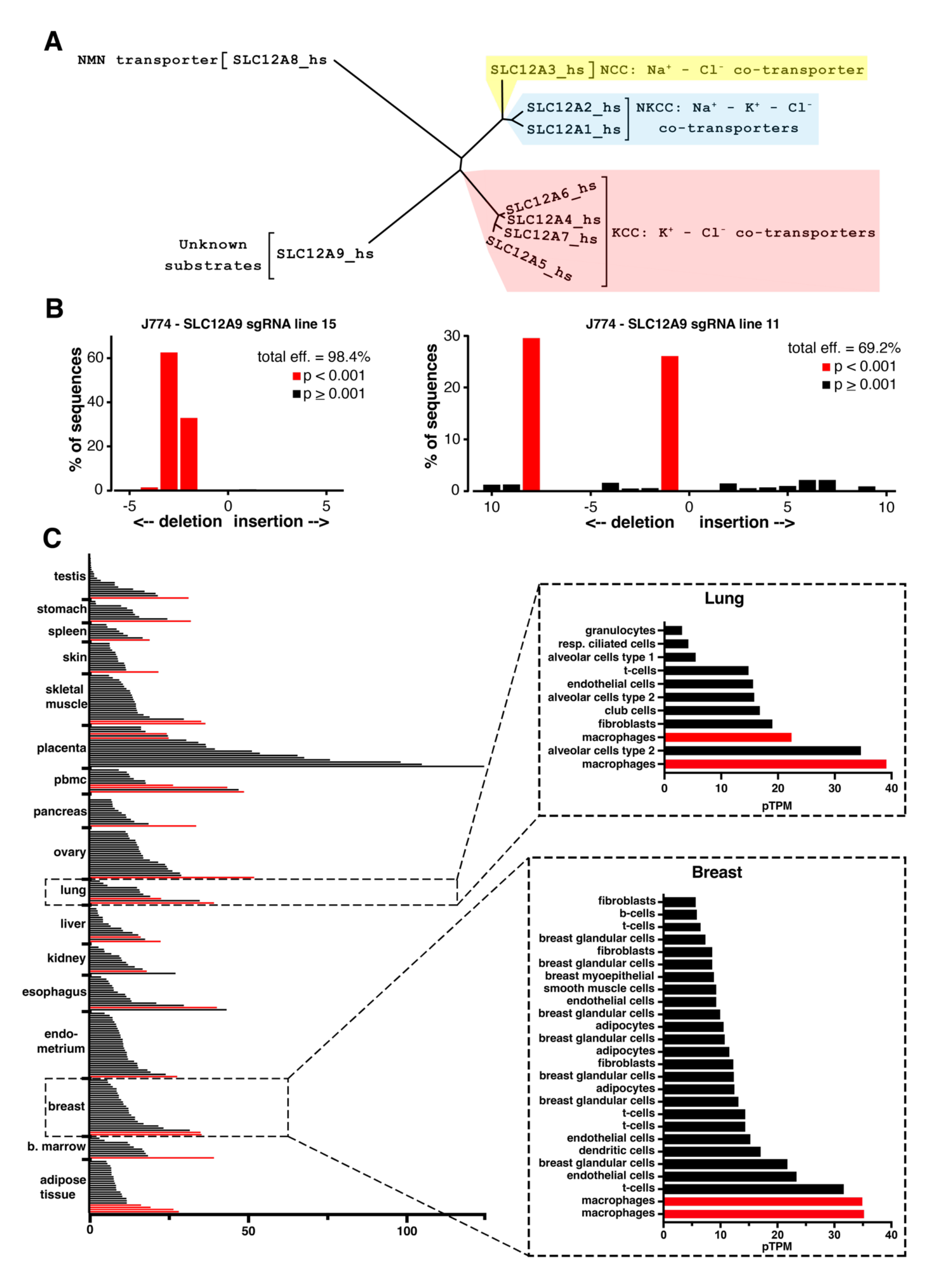
SLC12A9 an “orphan” member of the Cation – Chloride Co-transporter family. **A.** Phylogenetic homology tree of the CCC (SLC12A) family of solute carriers. Distances are inversely proportional to the degree of identity. The analysis was performed using the Phylogeny Analysis tool of *Methodes et Algorithmes pour la Bio-informatique*; LIRRM **B.** TIDE sequencing analysis of genomic DNA extracted from clonal Cas9-J774A.1 cell lines that had been transduced with an SLC12A9-targeting sgRNA. **C.** Cell-type transcript expression profile (protein transcripts per million) of SLC12A9 <u>47</u>; red bars highlight macrophages of each tissue.

**Figure S3.**
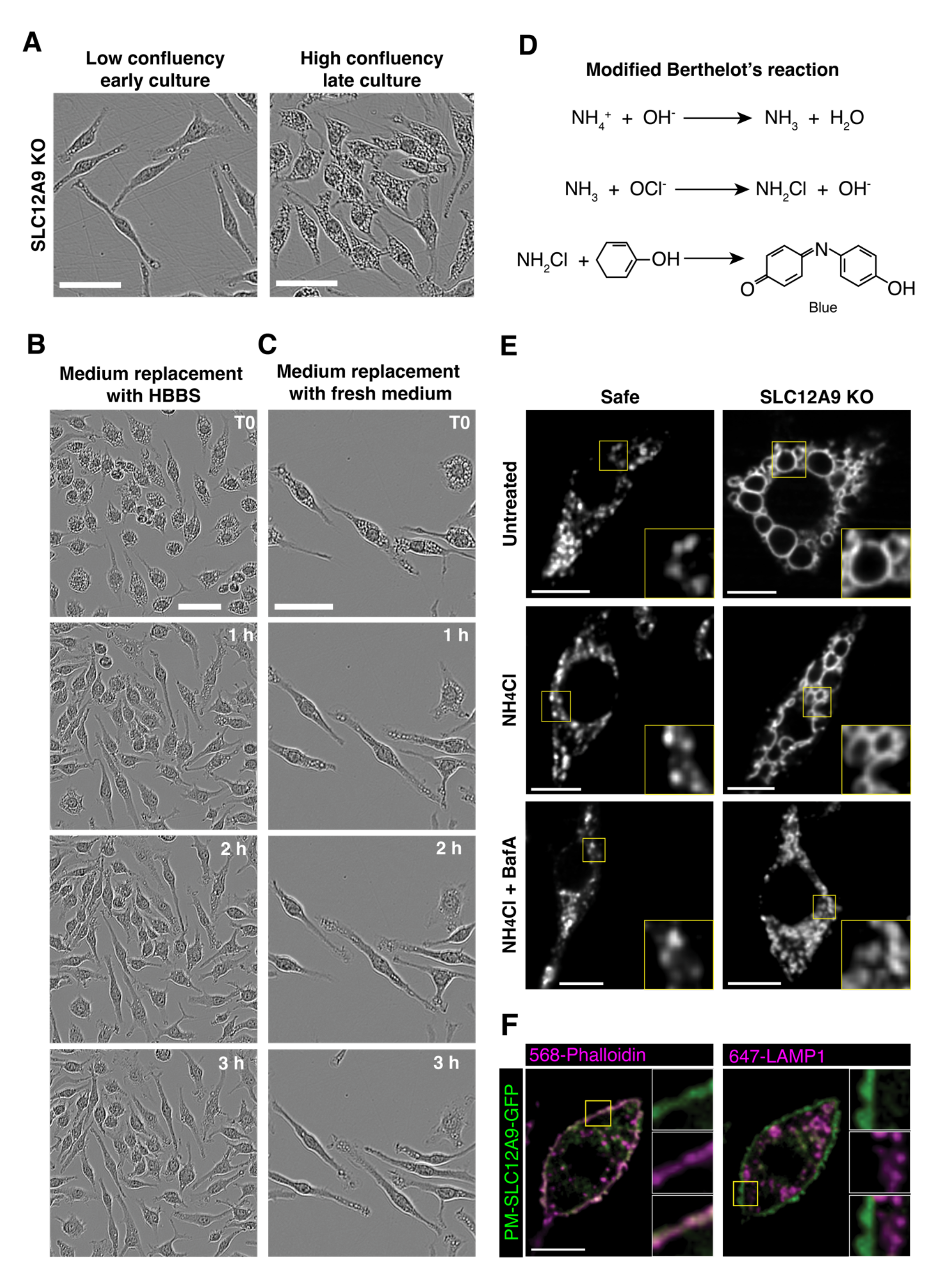
SLC12A9 transports ammonium across biological membranes. **A.** Brightfield Incucyte micrographs showing a low confluency, early culture of SLC12A9 KO J774A.1 cells (left) and a high confluency late culture of the same line (right). **B and C.** Brightfield Incucyte micrographs showing SLC12A9 KO J774A.1 cells at different time-points after culture medium replacement with (**B**) Hanks balanced salt solution or (**C**) fresh growth medium. **D.** Modified Berthelot’s reaction to measure total ammonia + ammonium levels. **E.** Representative confocal slices showing LAMP1 staining of control (top) and SLC12A9 KO (bottom) J774A.1 macrophages cultured under the described conditions. **F.** Representative confocal slices of J774A.1 macrophages expressing GFP-SLC12A9-LLY8/9/11AAA (green in both micrographs), stained with 568-phalloidin (magenta in the left micrograph) and with LAMP1 (magenta in the right micrograph). Scale bars in A., B., and C. = 20 µm. Scale bars in E., and F. = 10 µm.

**Figure S4.**
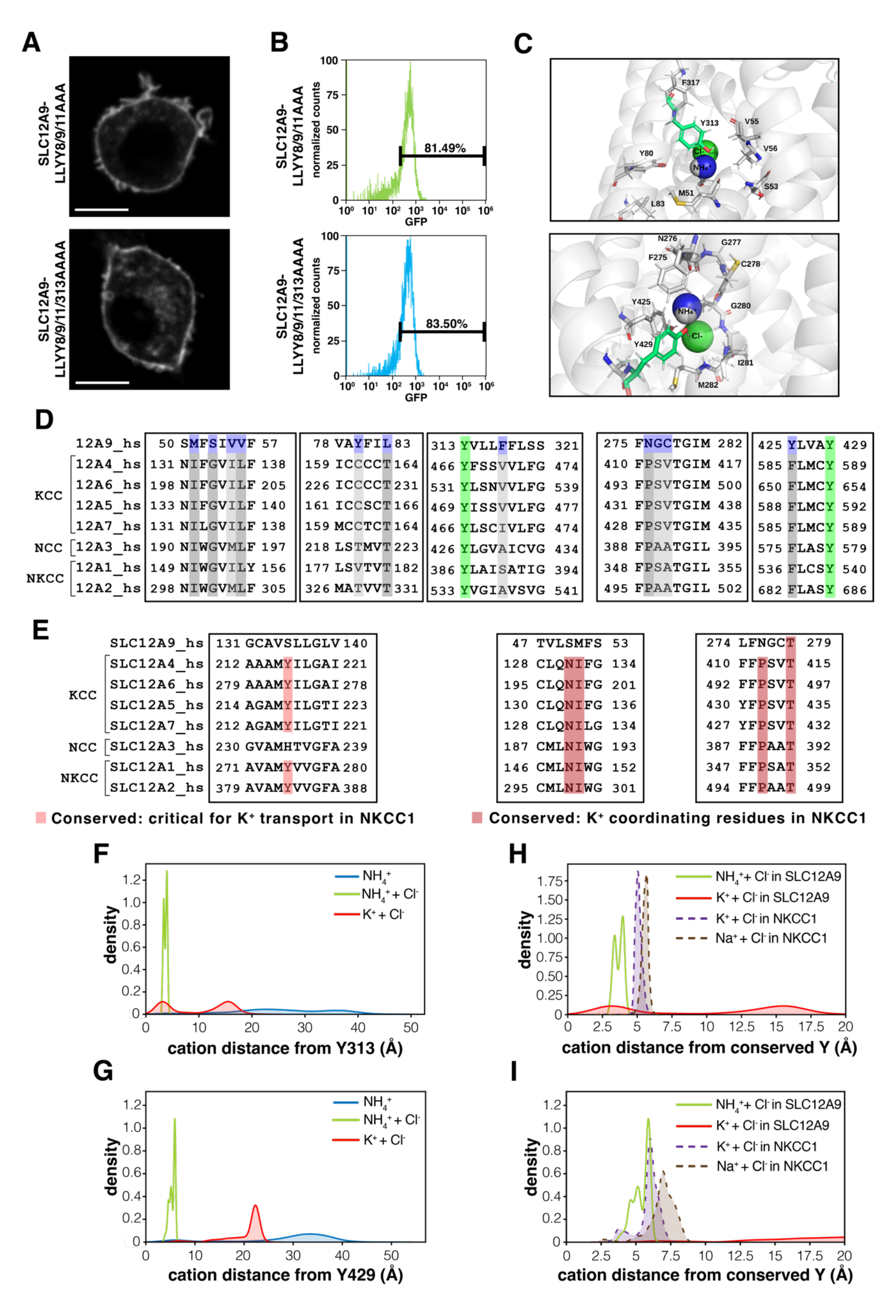
Ammonium transport by SLC12A9 is dependent on chloride binding. **A.** Representative confocal slices showing the subcellular localization of SLC12A9-LLY 8/9/11-GFP (top) and SLC12A9-LLYY 8/9/11/313 AAAA-GFP expressed in J774A.1 macrophages. **B.** Histogram showing the distribution of cell populations based on GFP fluorescence intensity of J774A.1 cells expressing SLC12A9-LLY 8/9/11-GFP (top; green curve) or SLC12A9-LLYY 8/9/11/313 AAAA-GFP (bottom; blue curve). **C.** Magnification of the SLC12A9-predicted structure (AlphaFold) showcasing the binding pockets centered on Y313 (Top) and Y429 (Bottom); highlighted are residues within 6 Å of the ions. **D.** Amino acid sequence conservation of members of the CCC (SLC12A) family; highlighted: in green: conserved tyrosines critical for transport in SLC12A2; in dark gray: residues that are conserved in all members except for SLC12A9 (blue); in light gray: residues that are partially conserved within the family but are not conserved in SLC12A9 (blue). **E.** Amino acid sequence conservation of members of the CCC family highlighted in red are conserved residues that are important for SLC12A2 transport of potassium. **F – I.** Histograms generated from simulations shown in movies x showing how long each of the specified cations spent at distances from residues Y313 **(F and H)** and Y429 **(G and I)** in SLC12A9 (AlphaFold predicted structure) and from residues Y533 **(H)** and Y686 **(I)** in SLC12A2 / NKCC1 (AlphaFold predicted structure). Scale bars = 10 µm.

**Figure S5.**
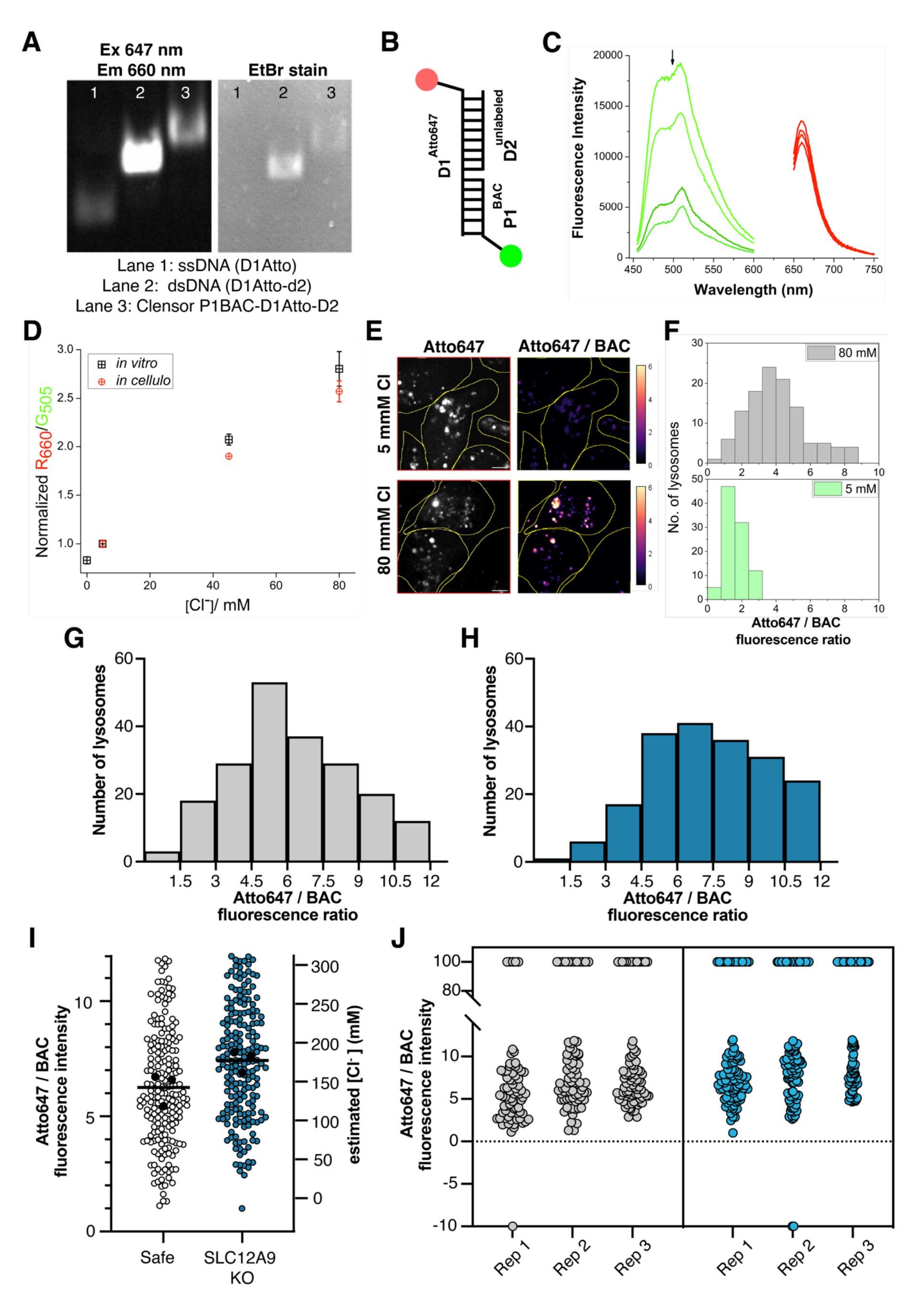
Synthesis, validation and use of Clensor. **A.** PAGE analysis of the synthesis of Clensor; lane 1 was loaded with Atto647-labeled single stranded DNA, lane 2 was loaded with Atto647-labeled double stranded (ds) DNA and lane 3 was loaded with Clensor. The left panel shows Atto647 fluorescence, and the right panel shows ethidium bromide signal. **B.** Schematic illustrating Clensor, a DNA-based ratiometric Cl-sensor: PNA strand P1 is the sensing domain containing BAC; DNA strand D1 is the ratiometric domain containing Atto647; DNA strand D2 is the organelle targeting domain. **C.** Emission spectra for BAC (green curves) acquired between 450 – 600 nm after excitation at 430 nm. Emission spectra for Atoo647 (red curves) acquired between 645 – 750 nm after excitation at 640 nm. **D.** Normalized ratios of red / green mean fluorescence intensities at different concentrations of chloride *in vitro* (gray and white squares) and *in cellulo* (red and white circles). **E.** Confocal micrographs showing J774A.1 cells after clamping with 5 mM (top) or 80 mM (bottom) of Cl-. Depicted are the Atto647 fluorescence (left) and a pseudo-colored representation of the red / green ratio (right). **F.** Histograms representing a typical spread of Red / Green ratios of lysosomes clamped at 5 and 80 mM of chloride (n ≥ 10 cells, ≥90 compartments). **G and H.** Histograms showing the number of lysosomes at different ranges of red / green fluorescence ratios in safe (**G**) and SLC12A9 KO cells (**H**). **I.** Values of red / green fluorescence (left Y axis) and estimated chloride concentration in mM (right Y axis) of individual lysosomes of safe and SLC12A9 KO cells loaded with Clensor. Highlighted as black horizontal are the mean values of the means (black data points) of three independent replicates per sample with at least 15 cells and ≥ 90 compartments. **J.** Values of red / green fluorescence of individual lysosomes of safe and SLC12A9 KO J774A.1 cells loaded with Clensor. Shown are values for each independent replicate.

**Figure S6.**
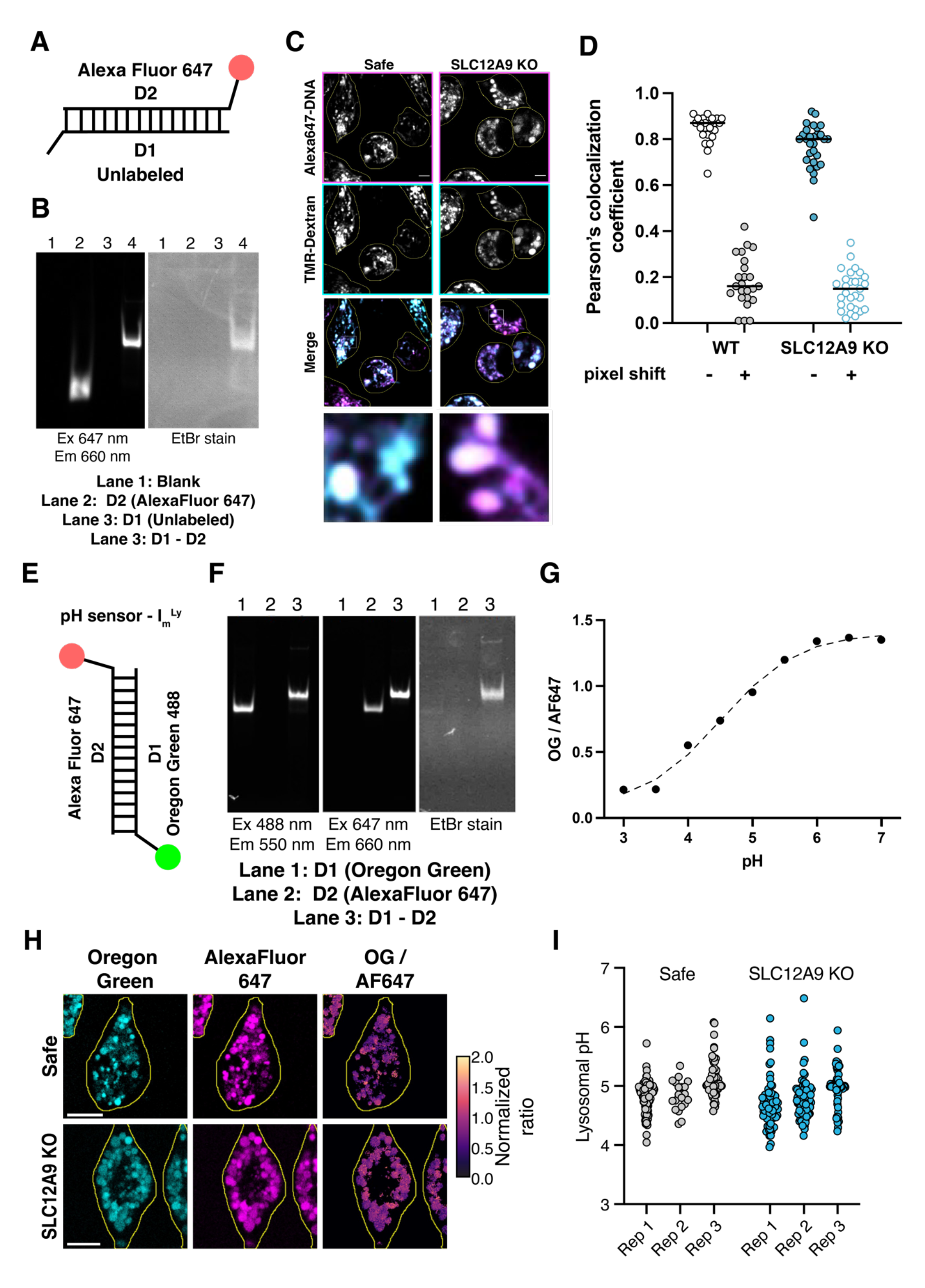
Synthesis, validation and use of I_m_^Ly^. **A.** Schematic illustrating a double stranded (ds) DNA-based AF647 probe; DNA stand 1 is unlabeled, and DNA strand 2 is tagged with AlexaFluor 647. **B.** PAGE analysis of the synthesis of the AF647-labeled DNA-based probe. Lane 1 was blank; lane 2 was loaded with AlexaFluor647-labeled single stranded DNA, D2; lane 3 was loaded with unlabeled single stranded DNA D1 and lane 3 was loaded with double stranded DNA duplex D1-D2. The left panel shows AlexaFluor647 fluorescence and the right panel shows ethidium bromide signal. **C.** Fluorescent micrographs of Safe (left) and SLC12A9 KO (right) J774A.1. cells pre-loaded with TMR dextran for lysosomal labeling and then loaded with an AF647-labeled ds DNA probe. Depicted are the AF647 fluorescence (top row), the TMR-dextran fluorescence (2nd row from the top), pseudo-colored merged representation of the AF647 and TMR channels (3rd row from the top) and a magnification of a region of interest from the merged representation (bottom row). **D.** Co-localization analysis of the experiments described in **C** showing the analysis with a pixel shift as a control. **E.** Schematic illustrating a double stranded (ds) DNA-based ratiometric pH sensor I_m_^Ly^; DNA stand 1 is tagged with Oregon Green, and DNA strand 2 is tagged with AlexaFluor 647. **F.** PAGE analysis of the synthesis of the pH sensor ImLy. Lane 1 was loaded with Oregon-Green-labeled single stranded DNA, D1; lane 2 was loaded with AlexaFluor647-labeled single stranded DNA D2; and lane 3 was loaded with double stranded DNA duplex D1-D2. The left panel shows the Oregon Green fluorescence, the middle panel shows the AlexaFluor647 fluorescence, and the right panel shows ethidium bromide signal. **G.** Normalized ratio of Oregon Green and AlexaFluor647 fluorescence intensities within I_m_^LY^ pH sensor at pH values ranging from 3 – 7; Oregon Green was excited with a 488 nm laser and an emission spectrum was acquired from 500 to 650 nm; AlexaFluor 647 was excited with a 647 nm laser and an emission spectrum was acquired from 660 to 750 nm. **H.** Representative confocal slice of a J774A.1 macrophages loaded with I_m_^Ly^ pH sensor. Shown are safe (top) and SLC12A9 KO (bottom) cells. Depicted are pseudo-colored images of the Oregon green fluorescence (left), AF647 fluorescence (middle) and a representation of the ratio of Oregon Green and AF647 fluorescence (right). **I.** Calculated pH values of individual lysosomes of cells loaded with I_m_^Ly^ pH sensor. Shown are the values of each independent replicate experiment in which at least 120 lysosomes were quantified from at least 10 cells.

